# Cold-passaged isolates and bat-swine influenza A chimeric viruses as modified live-attenuated vaccines against influenza A viruses in pigs

**DOI:** 10.1101/2022.06.20.496807

**Authors:** Annika Graaf, Philipp P. Petric, Julia Sehl-Ewert, Dinah Henritzi, Angele Breithaupt, Jacqueline King, Anne Pohlmann, Fabian Deutskens, Martin Beer, Martin Schwemmle, Timm Harder

**Author notes:** Correspondence: Prof. Dr. Timm Harder.

## Abstract

Swine influenza A virus (swIAV) infections in pig populations cause considerable morbidity and economic losses. Frequent reverse zoonotic incursions of human IAV boost reassortment opportunities with authentic porcine and avian-like IAV in swine herds potentially enhancing zoonotic and even pre-pandemic potential. Vaccination using adjuvanted inactivated full virus vaccines is frequently used in attempting control of swIAV infections. Accelerated antigenic drift of swIAV in large swine holdings and interference of maternal antibodies with vaccine in piglets can compromise these efforts. Potentially more efficacious modified live-attenuated vaccines (MLVs) bear the risk of reversion of MLV to virulence. Here we evaluated new MLV candidates based on cold-passaged swIAV or on reassortment-incompetent bat-IAV-swIAV chimeric viruses. Serial cold-passaging of various swIAV subtypes did not yield unambiguously temperature-sensitive mutants although safety studies in mice and pigs suggested some degree of attenuation. Chimeric bat-swIAV expressing the hemagglutinin and neuraminidase of an avian-like H1N1, in contrast, proved to be safe in mice and pigs, and a single nasal inoculation induced protective immunity against homologous challenge in pigs. Reassortant-incompetent chimeric bat-swIAV vaccines could aid in reducing the amount of swIAV circulating in pig populations, thereby increasing animal welfare, limiting economic losses and lowering the risk of zoonotic swIAV transmission.

## Introduction

Worldwide, domestic pigs are clinically affected by swine influenza A virus (swIAV) infections comprising the hemagglutinin (HA) subtypes H1 or H3, and the neuraminidase (NA) subtypes N1 or N2. Within each subtype several viral lineages exist which are defined on basis of their original host origin as avian (av), human (hu) or pandemic (pdm). A corresponding nomenclature proposed by Anderson et al. (2016) related to phylogenetic analyses refers to those lineages as 1A (=pdm), 1B (=hu), and 1C (=av). Infections are associated with disease and economically significant losses, especially in piglet rearing and reproduction (Brown, 2000; Rajao et al., 2014; Vincent et al., 2014). Complex and synergistic interactions of swIAV infections with other pathogenic or opportunistic viral and bacterial infectious agents contribute to an aggravation of disease which is broadly referred to as the porcine respiratory disease complex, PRDC (Brockmeier, 2002; Deblanc et al., 2017; Pirolo et al., 2021; Rajao et al., 2018; Saade et al., 2020). Removing pathogens from the PRDC, e.g. by antibiotic treatment or vaccination, disentangles synergistic effects resulting in an overall improved animal health status and in reduced economic losses (Fachinger et al., 2008).

SwIAV are closely related to IAV circulating in the human population (Anderson, 2021; Chauhan et al., 2020). In fact, reciprocal transmissions across the porcine/human interface are observed regularly and significantly contribute to the expansion of swIAV variability in pig populations (Chastagner et al., 2019; Nelson et al., 2012; Neumann et al., 2009). Pigs are also susceptible to IAV of avian origin, further adding to the diversity of circulating swIAV. Pigs therefore resemble transitional influenza virus hosts or “mixing vessels” which facilitate the reassortment of IAV genomes of different host origins (Hass et al., 2011; Ma et al., 2009; Webster, 1997). This opens wide opportunities for zoonotic transmissions of reassorted swIAV with unknown phenotypic properties which might include pandemic potential as documented by the most recent human influenza pandemic of 2009 originating from reassorted swIAV in Mesoamerica (Short et al., 2015).

Recent extensive studies specified a broad and complex reservoir of swIAV with zoonotic and even prepandemic potential in European, Asian and American domestic swine populations (Henritzi et al., 2020; Sun et al., 2020; Vincent et al., 2008). In European pig populations, a dramatically increased genetic complexity of four lineages of swIAV, H1N1 of avian and of pandemic origin, H1N2 and H3N2, various reassortants thereof and further sporadic human-to-swine spillover variants has been described since 2010 (Henritzi et al., 2020; Krog et al., 2017). Increasing sizes of swine breeding herds and other intensification measures of pig production have provided swIAV with enriched opportunities to establish enzootic, self-sustaining infections in large herds (Pitzer et al., 2016; Ryt-Hansen et al., 2019). Swine influenza, in Europe, is currently not subject to any statutory reporting, and harmonized, officially orchestrated monitoring programs and control strategies are not implemented.

Besides improving biosafety to block swIAV incursions into swine herds, vaccination is the key preventive measure to combat influenza in pigs (Rahn, 2015; Sandbulte et al., 2015; Vincent, 2017). Currently available commercial influenza vaccines for use in swine are essentially based on adjuvanted inactivated whole virus (AIWV) vaccines. Administration of oil-adjuvanted AIWV by parenteral injection establishes a short-lived systemic immunity (Van Reeth et al., 2013) comprising strain-specific neutralizing antibodies primarily against the HA and to a lesser extent against the NA glycoproteins of swIAV as the main effectors conferring protection (Padilla-Quirarte et al., 2019). Cell-mediated immunity induced by AIWV has not been investigated thoroughly due to the complexity of the assays and because of a general consent on the predominant role of neutralizing antibodies. The latter often confer a narrow protection window and cross-protection against a wide spectrum of antigenically distinct variants is rare. Frequent re-vaccinations are not always effective to counteract these shortcomings; instead, complex cross boostering scheme using carefully selected heterologous vaccine antigens can be successful in widening protection efficacy (Chepkwony et al., 2020). In addition, transfer of colostral maternal immunity from vaccinated sows to suckling piglets does not induce sterile immunity (Genzow et al., 2018): While clinical sequelae of an swIAV infection are reduced in such piglets, they virus infection itself is not prevented so they have been suspected drivers of endemic swIAV infections in large herds (Ryt-Hansen et al., 2020). At the same time, interference with maternally derived swIAV antibodies restricts the use of AIWV in piglets during their first ten weeks of life, and most vaccines are licensed for use in pigs from 56 days of age only (Deblanc et al., 2018; Everett, 2021).

Modified live-attenuated vaccines (MLVs), in contrast, are known to induce both humoral and T-cell mediated immunity including a mucosal component depending on the application route (Pollard et al., 2021). Mucosally administered replication-competent influenza vaccines therefore are expected to combine several advantages: they supposed to induce immunity in the airway epithelium, i.e. the primary location of viral entry and initial replication and this abrogates intraepithelial spread, and decreased viral excretion (Lavelle et al., 2021). In addition, mucosal application to suckling piglets would evade interference with maternal derived, systemic (non-mucosal) neutralizing antibodies. Several attempts to develop efficacious influenza MLV for use in swine have been reported, culminating in the licensing of a bivalent MLV which was attenuated by a C-terminal deletion of the NS protein (Genzow et al., 2018; Solorzano et al., 2005; Vincent et al., 2007). Despite initial success, this vaccine had to be withdrawn from the market after a short period of use in North American swine populations due to reassortment with endemic wild-type swIAV and, hence, indirect reversion to virulence (Sharma et al., 2020). Thus, despite of their obvious advantages, MLV vaccines can bear significant risks for reversion to virulence and disease, and also the potential for shedding which, in the case of swIAV, created public health concerns (Vincent, 2017).

Temperature-sensitive (*ts*) mutant IAV realize another attenuation principle that has been utilized in MLVs for human use: they show an at least 100-fold reduced replication efficacy at higher incubation temperatures *in vitro*; hence, they replicate efficiently in upper airway epithelia but not at higher temperatures in deeper airway tissues (Broadbent et al., 2014; Chan et al., 2008; Chen et al., 2010; He et al., 2013; Isakova-Sivak et al., 2011). While attenuated (*att*), *ts* mutants remain competent to induce protective mucosal immunity. Along this line, quadrivalent influenza vaccines have been approved for safe use in humans, particularly in children (Belshe, 2004). Nevertheless, such mutants still bear the intrinsic risk of reassorting with field viruses which challenges their safety. However, reassortment competence with circulating field viruses are blocked in reverse genetically (*rg*) constructed chimeric (*ch*) viruses carrying the HA and NA of mammalian or avian IAV in the backbone of bat influenza A viruses (bt IAV) (Ciminski et al., 2017; Juozapaitis et al., 2014; Ma et al., 2015; Schon et al., 2020; Yang et al., 2017; Zhou et al., 2014). This would confer a substantial gain in influenza MLV safety.

In this study, safety and efficacy of MLV vaccines based on newly developed, serially cold-passaged swIAV of several European lineages or on ch-bt IAV were assessed in mice and pigs. Evidence from single vaccination-challenge experiments in pigs using ch-bt IAV showed clinical protection and reduced excretion and spread of challenge virus.

## Results

### Serial cold-passaging of swIAV at 28°C resulted in *ts-like* mutant influenza viruses

In order to obtain a sufficiently high-titred virus stock for serial passaging experiments, selected parental swIAV 531 (H3N2), 541 (H1pdmN2) and 1670 (H1avN1), all obtained from pigs with respiratory disease on European farms (Table 1), had received 2-4 passages in MDCK-II cells at 37°C. As expected, these low passage isolates, designated par531, par541 and par1670 generated higher virus titers in MDCK-II cell culture at 37°C compared to 28°C (Figure 1 A, D, G). Maximum titers at 37°C were reached within 24 hours and measured up to three log_10_ steps higher compared to 28°C until 72 hours. Next, 60 serial cold passages (cp) were performed with the parental swIAV in either MDCK-II or in a swine testicle cell line (ST-0606) resulting in the virus stocks cp531 and cp541 and cp1670. Only cp1670 passaged in MDCK-II cells but not in ST cells produced higher virus titers at an earlier timepoint at 28 versus 37°C with maximum titer of up to 10^6^ TCID_50_/mL (Figure 1 F, I). These growth differences were less striking for cp531 and cp541 passaged on both MDCK-II and ST cells (Figure 1 D-G, E-H). However, all cold passaged viruses induced a cytopathic effect at 28°C but not at 37°C (not shown). Thus, only cp1670 revealed replication features described for *ts* mutant influenza viruses (Martinez-Sobrido et al., 2018).

**Figure 1.**
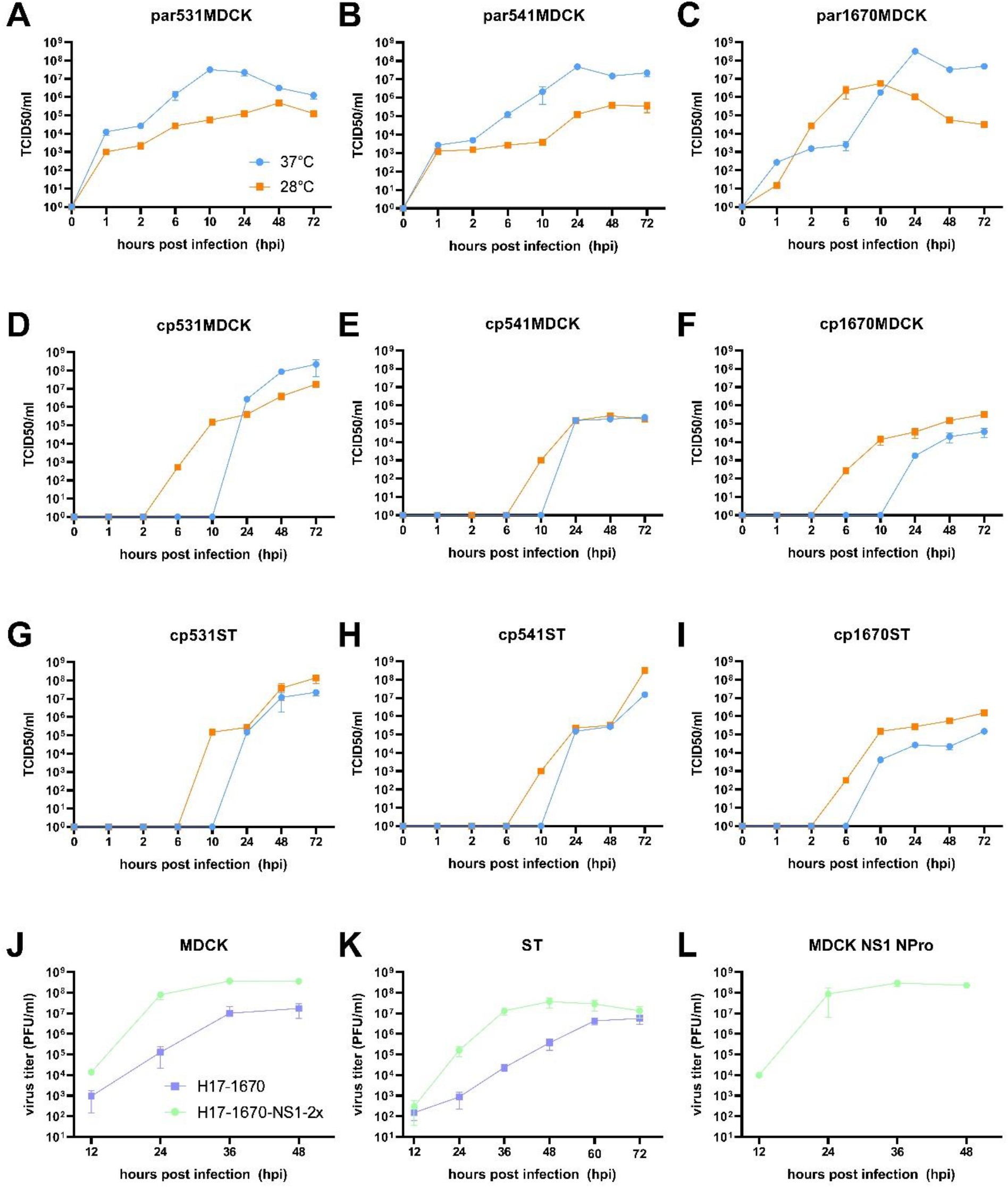
Growth kinetics of parental swIA viruses (A-C), cold-passaged isolates thereof after 60 serial passages in (D-F) MDCK-II or (G-I) ST-0606 cell culture at 37°C (blue) versus 28°C (orange) and growth kinetics on (J) MDCK-II, (K) ST-0606 and (L) MDCK-NS1-NPro cell cultures infected with ch-bt IAV candidates H17-1670 (purple) and H17-1670-NS1-2x (green). Mean values were calculated for each virus from three independent biological replicates.

**Table 1.** Characteristics of selected virus isolates and strains used for serial cold passaging in cell culture, for generation of chimeric-bt IAV by reverse genetics (*rg*) and for inoculation and challenge experiments in mice and pigs. *MDCK-II – Madin Darby canine kidney cell; ST-swine testicle cell; nd – not defined ** Reversion to cytopathic virus replication after ten blind passages at 37°

| Isolate-ID | Subtype | Parental virus | Cell line | Passage | Vaccine candidate | Reversion to thermophilic growth ** |
| --- | --- | --- | --- | --- | --- | --- |
| A/swine/Netherlands/AR531/2015 | H3N2 | par531 | MDCK-II | 60 | cp531 | - |
|  |  |  | ST |  |  | - |
| A/swine/Germany/R541/2012 | H1pdmN2 | par541 | MDCK-II | 60 | cp541 | + |
|  |  |  | ST |  |  | + |
| A/swine/Germany/AR1670/2014 | H1avN1 | par1670 | MDCK-II | 60 | cp1670MDCK<br>cp 1670ST | + |
|  |  |  | ST |  |  | + |
| A/swine/Germany/AR1670/2014 | H1avN1<br>+ H17N10 | par1670 | MDCK-II | rg | H17-1670 | nd |
| A/little yellow<br>shouldered bat/Guatemala/164/2009 |  |  | MDCK-NS1-Npro | rg | H17-1670-NS1-2x | nd |

### The cold-adapted swIAV reversed to cytopathogenic replication following consecutive passaging at 37°C

Whole genome sequencing by the MinION technologies revealed for the majority of the cp swIAV, non-synonymous mutations were mainly detected in gene segments coding for the polymerase complex (PA, PB1, PB2 and NP). In contrast, hardly any coding mutations occurred in the HA, NA, M and NS segments (Table 2). The parallel passages of the same parental isolate in MDCK-II and ST cell lines, respectively, resulted in concordant amino acid substitutions at positions PB2 D87G and E158G, in cp531, and cp1670 (highlighted in bold type in Table 2). Illumina whole genome sequencing confirmed their presence as did Sanger sequencing of short PCR amplicons generated with specific primers for the relevant regions (not shown). None of these mutations have previously been described in *ts* mutants of other other IAV. However, Zhou et al. (2011) showed that PB2-E158G might play a role in the adaptation of avian PB2 genes to other mammals. No coding mutations occurred in the regions of the HA and NA proteins known as epitopes of humoral immunity (Petukhova et al., 2017). Based on these findings, the serially cold-adapted viruses were considered antigenically similar to their parental isolates.

**Table 2.**
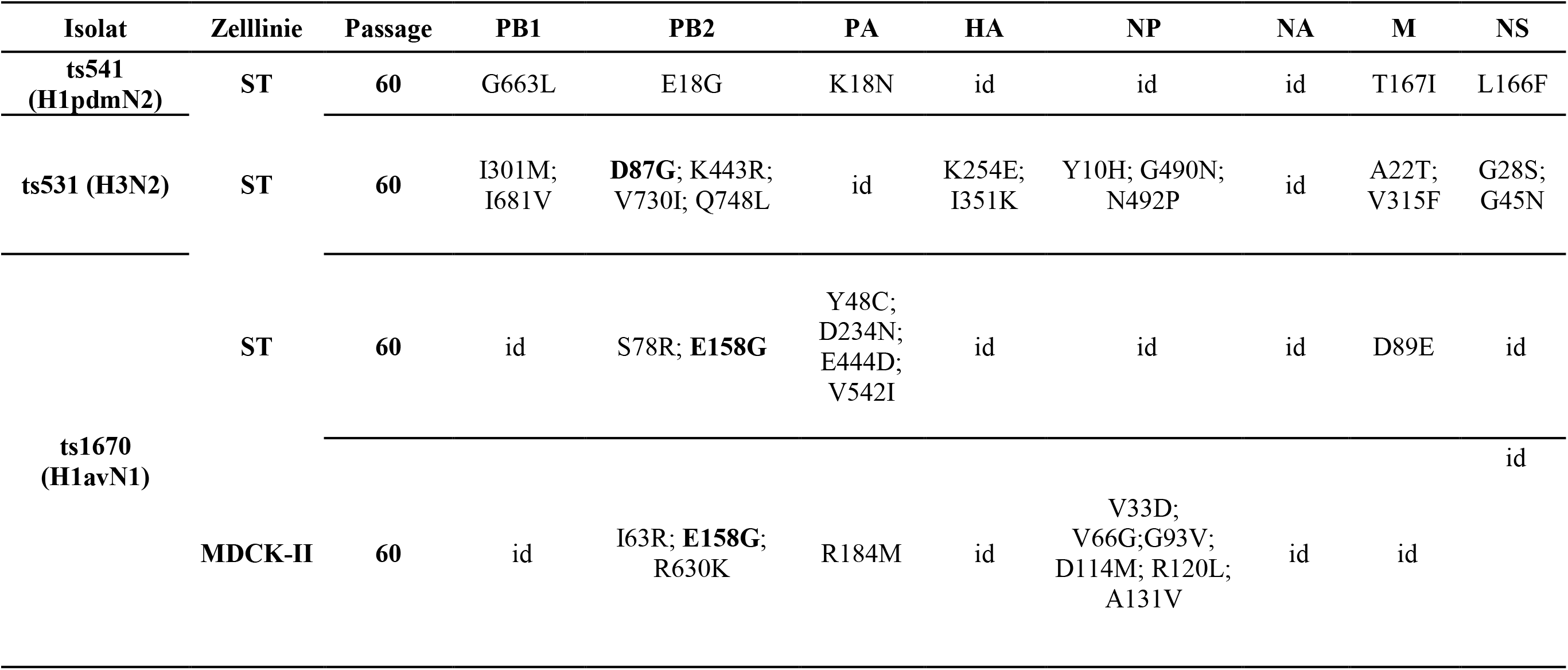
Non-synonymous mutations of serially cold-passaged swIAV isolates 531 (H3N2), 541 (H1pdmN2), and 1670 (H1avN1) of passage 60 in ST or MDCK-II cell cultures compared to the corresponding low passage parental virus isolate replicated in MDCK-II cells at 37°C. Colors indicate parallel arising amino acid substitutions after cold passaging in both ST and MDCK-II cell cultures. *id-identical to the parental virus

Next, each virus, after 60 cold passages, was subjected to ten serial passages in both MDCK-II and ST cells at 37°C as shown in Figure 2 to provide information on the frequency of possible reversion to the thermophilic cytopathic replication. While cp531, cp541 and cp1670 initially did not induce cytopathic effects while passaged at 37°C, return of cytopathogenic replication at 37°C was noticed in all strains with the exception of cp531 (H3N2) (supplemental table 1). To determine the genetic basis of the phenotypic reversion, we analyzed the mutations within PB2 segment using an NGS approach. Interestingly, cp541 (H1pdmN2) showed a reversion of its mutation E158G and cp531 reversed mutations V730I and Q748L, whereas cp1670 showed no reversions of the mutations in PB2 segment selected during cold-passaging. In summary, cp541 and cp1670 reversed to cytopathogenic replication following consecutive passaging at 37°C accompanied by reversion of the mutation obtained during passaging at 28°C or *de novo* mutations.

**Figure 2.**
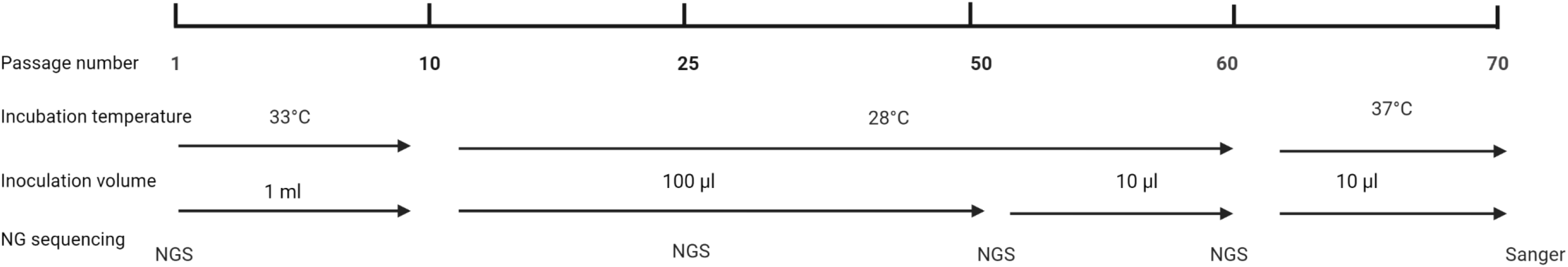
Scheme of the *in vitro* generation of serially cold passaged swine Influenza A viruses.

### Ch-bt IAV encoding wildtype or a NS1 with IFN antagonistic are replication competent

In addition to the cold-adapted swine IAV, reassortment-incompetent vaccine candidates comprising recombinant ch-bt IAV expressing the 1670 glycoproteins (H17-1670) and, as an additional safety measurement, a ch-bt NS1 double-mutant virus encoding the NS1 R39A and K42A mutations (H17-1670-NS1-2x) were generated (Figure 3). To analyze their replication competence, viral growth kinetics were performed on MDCK-II and ST cells (Figure 1 J-L). The H17-1670 virus showed efficient growth on both cell lines, reaching titers of up to 8 log_10_ PFU/ml. Compared to that, the H17-1670-NS1-2x virus showed highly impaired replication efficiency on both cell lines, with viral titers being up to 2 log_10_ lower than for the NS-wt virus at most timepoints. Interestingly, both viruses reach similar titers on ST cells after 72 hpi. As the two NS1 mutations were described to impair the IFN-antagonist function of the virally encoded NS1 protein, additional growth kinetics were performed on MDCK-NS1-NPro cells, expressing functional IFN-antagonists of an IAV strain (NS1) and bovine viral diarrhea virus (NPro) (Figure 1L). Consistent with our previous observation with ch-bt NS1 double-mutant virus expressing the H7 and N7 of A/SC35M (Juozapaitis et al., 2014), on this cell line, the H17-1670-NS1-2x virus showed identical replication efficiency as the wt NS virus on the MDCKII cells (compare Figure J, L). Of note, viral titers of both ch-bt IAVs are sufficiently high to be used for *in vivo* experiments.

**Figure 3.**
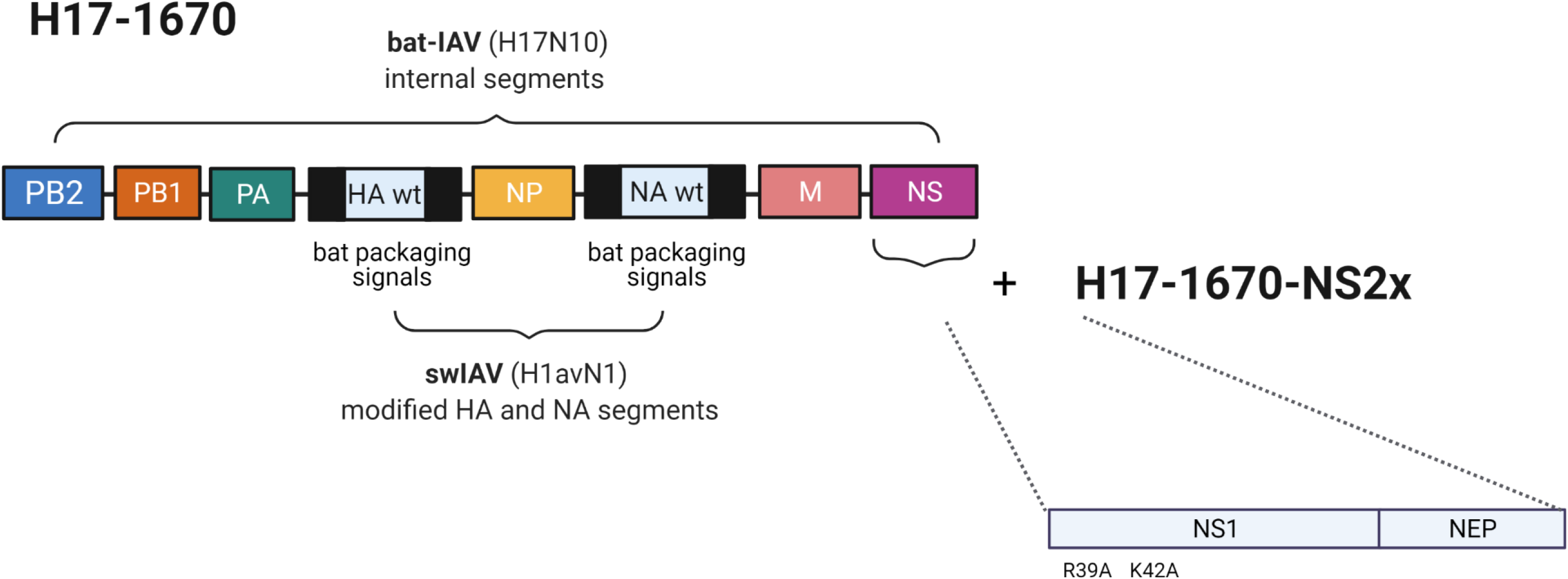
Genome organization of chimeric-bat-swine (HA, NA) influenza A viruses.

### Swine adapted IAV and bat viruses show different degrees of attenuation in mice

Virus strain A/Puerto Rico/8/1934 (H1N1) [PR8] was used as a virulent positive control to weigh clinical signs observed with parental and cp swIAV as well as ch-bt viruses. A predominantly severe clinical course associated with pneumonic alterations in the lung became evident when an intranasal inoculum volume of 40 µL of PR8 was used (supplemental figure 1). Four out of 8 inoculated mice reached humane endpoints within 10 days of observation. All inoculated mice showed higher RNA viral loads in lung tissues than in trachea or nose. A necrotizing bronchointerstitial pneumonia was diagnosed with abundant, intralesional IAV antigen in bronchiolar and alveolar epithelial cells of inoculated animals at 2 and 10 dpi (supplemental figure 2 A, B; supplemental figure 3E, J, O, T). Seroconversion occurred in all inoculated animals surviving until 10 dpi (supplemental figure 4). However, transmission to in-contact animals was not observed as none of the sentinels developed clinical symptoms nor showed any viral RNA loads by RT-qPCR examination in tissue samples and did not seroconvert.

None of the parental swIAV (par531, par541, par1670) induced clinical respiratory manifestations in the inoculated or in the contact mice. Substantial body weight reductions of up to 25% were measured in only one inoculated animal of group par541 and in three inoculated animals of group par1670 (supplemental figure 1). The mouse of group par541 recovered and survived until 10 dpi. Two mice of group par1670 showed significant weight loss but did not reach the 25% threshold until 10 dpi.

Compared to PR8 the virulence of par swIAV in mice was markedly reduced. Viral loads in two mice sacrificed per group at 2 dpi were higher in lungs compared to nasal mucosa. In the lungs, bronchointerstitial pneumonia was found in the majority of animals, again associated with IAV antigen detected in many bronchial and fewer alveolar epithelial cells (supplemental figure 2A). At day 10 pi, residual virus RNA was detected predominantly in lung tissues. This includes some contact animals suggesting transmission of parental swIAV by direct contact. Only a few mice in each par swIAV group seroconverted (supplemental figure 4). Accompanying this, bronchointerstitial pneumonia was found in several but not all animals, however viral antigen was still detectable in two mice after 1670MDCK infection (supplemental figure 2A; exemplarily shown in supplemental figure 3H).

Mice infected with either one of the four serially cold-passaged swIAV (cp531, cp541, cp1670MDCK, cp1670ST) as well as the respective contact animals did not show any respiratory signs of disease. However, two mice inoculated with cp1670MDCK showed sunken flanks, slightly ruffled fur associated with weight loss of < 25% between 4 to 7 dpi (supplemental figure 1). Viral loads were detected in turbinate tissues of cp inoculated mice at 2 dpi (Figure 4). Interestingly, for the two cp1670 viruses these were higher compared to par1670. Conversely, viral loads of cp swIAV in lung tissue at day 2 were markedly lower compared to par swIAV. Nevertheless, lesion-associated viral antigen was found in one mouse each after infection with cp531 or cp1670MDCK, respectively, but not after cp541 or cp1670ST infection (supplemental figure 2B).

**Figure 4.**
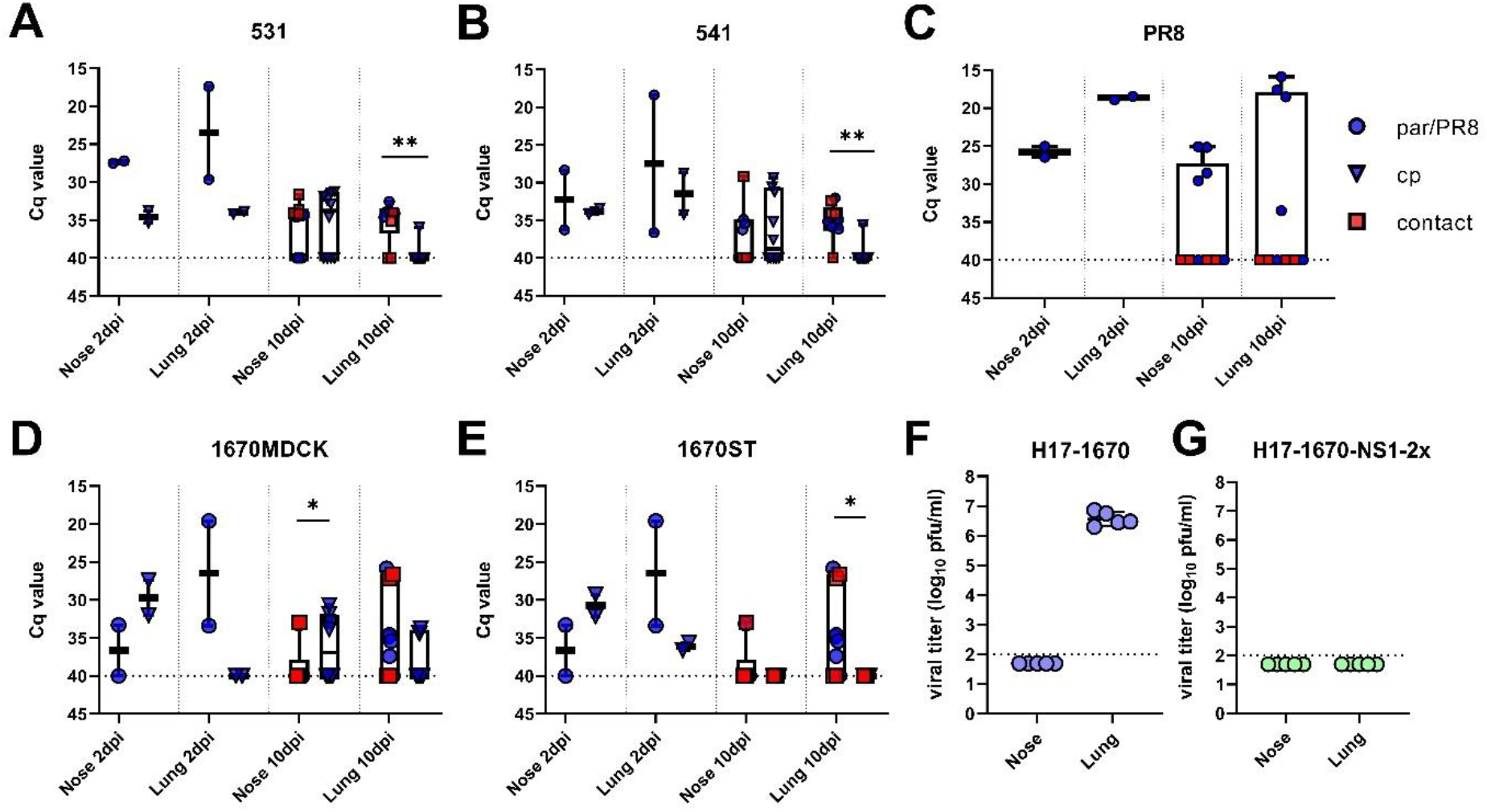
**(A-E)** Viral loads measured by RT-qPCR in nasal and lung tissues after inoculation of the different parental (par, rounded blue dots), cold-passaged (cp, blue triangles) swIAV or positive control (PR8) in C57BL/6 mice. Red squares indicated non-inoculated contact animals. (F-G) Viral titers measured by plaque assays in nasal and lung tissues after inoculation of C57BL/6 mice with ch-bt IAV candidates H17-1670 (purple) and H17-1670-NS1-2x (blue). *-*** indicate significance levels (p<0.05; <0.01; <0.001)

At 10 dpi, residual viral RNA was detectable in nasal tissues but hardly in lung tissues of cp inoculated mice. Within 10 days, seroconversion could be measured in some cp-inoculated mice but not in the contact animals, similar to the groups of the parental viruses (supplemental figure 4). None of the mice exhibited viral antigen in the lungs. However, in accordance with antigen labeling data from 2 dpi, infection with cp531 and cp1670MDCK led to pneumonia in all mice, indicating prior virus replication in lung tissue (supplemental figure 2A).

To evaluate the growth properties and pathogenicity of the ch-bt IAV vaccine candidates *in vivo*, five C57BL/6 mice each were intranasally inoculated with 10^4^ PFU/ml of either H17-1670 or H17-1670-NS1-2x. The mice were sacrificed 3 dpi to determine viral lung and nose titers (Figure 4 F-G). While the H17-1670 virus reached viral titers of up to 10^7^ PFU/ml in the lungs of infected mice, no virus could be isolated from the upper airway (nose). Interestingly, no infectious H17-1670-NS1-2x was identified in both the upper airway and lungs of infected mice, suggesting that loss of the RNA-binding activity in NS1 caused severe attenuation *in vivo*.

In summary, as expected, cp swIAV showed a different replication pattern in C57BL/6 mice compared to par swIAV. The predominant replication in nasal versus lung tissues suggests features of an *att* phenotype of cp swIAV in mice as described for *ts* mutant IAV (Broadbent et al., 2014; Isakova-Sivak et al., 2011). In addition, the level of cp swIAV replication in mice might be overall reduced compared to the parental viruses as less of the inoculated mice seroconverted. However, diagnosis of pneumonia after cp531 and cp1670MDCK infection indicates pulmonary replication of cp swIAV to a certain extent. In contrast, and similar to other ch-bt IAV (Juozapaitis et al., 2014), H17-1670 did not show obvious signs of disease despite efficient replication in the lung.

### Cold-passaged swIAV show varying degrees of attenuation in pigs

In order to analyze the safety of cp swIAV in pigs, eight pigs were intranasally inoculated with 10^6^ TCID_50_ of cp531, and at day 1 p.i. two contact pigs were associated. Results were compared to a group of four age-matched pigs which received the same dose of par531. Neither the par nor the cp virus induced clinical signs including a rise in body temperature, the exception being a single cp531-inoculated animal that showed a light cough at 3 dpi (supplemental figure 5A, B). Animals of both groups, including the cp sentinels, excreted virus nasally at days 2, 4 and 5 p.i. (Figure 5A, sentinels not shown). With regard to the dynamics and intensity of nasal excretion, there were no significant differences between the cp and par groups. Respiratory tissues of both groups were found to be virus positive at 4 dpi (Figure 5B). A significantly increased viral load of the parental virus was found in the bronchial swabs and in the BALF materials, whereas no significant differences were found for any of the remaining tissues. As expected, within the four to five days observation period, none of the animals seroconverted (not shown).

**Figure 5.**
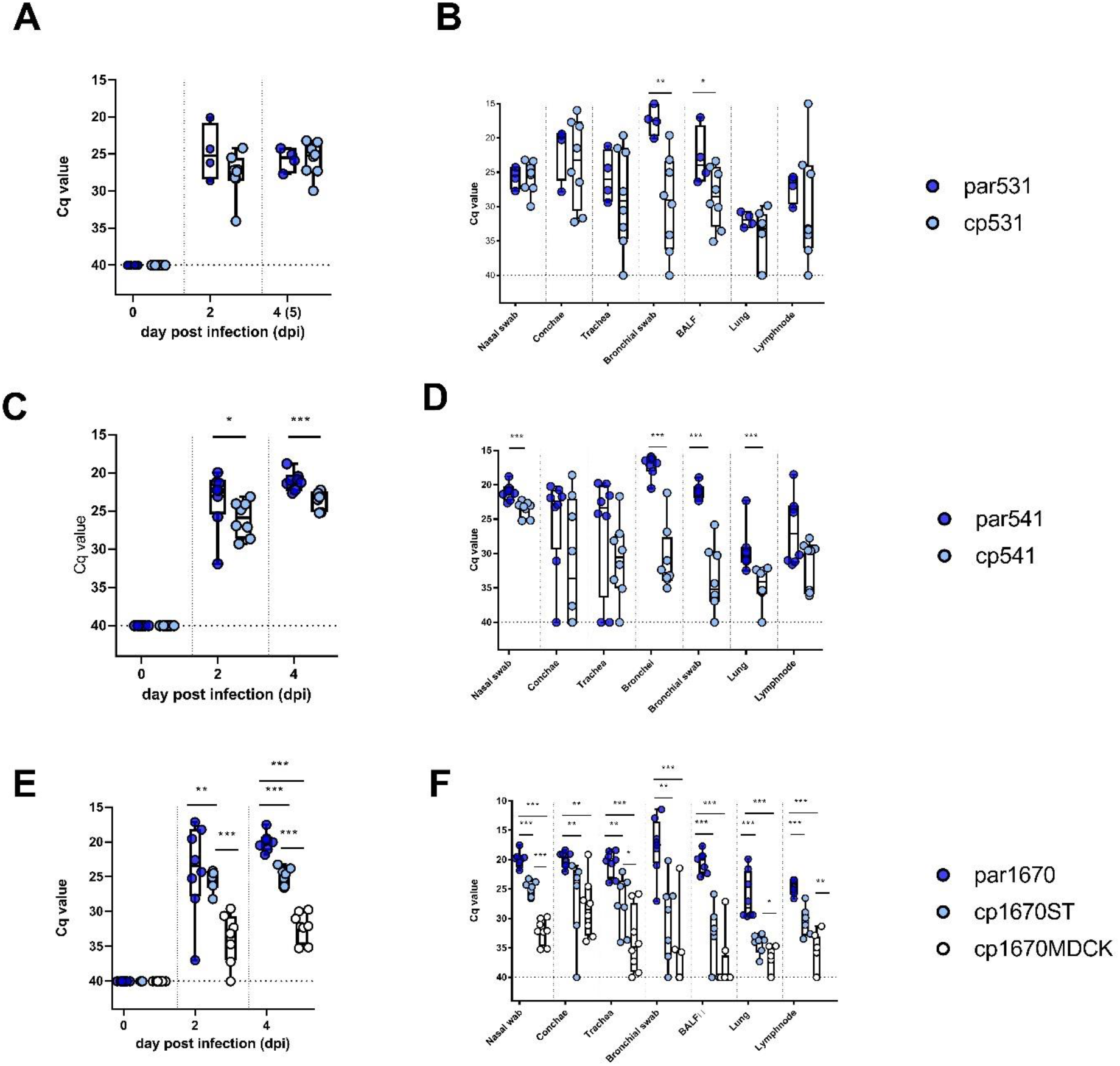
Nasal viral excretion (A, C, E) and viral loads in respiratory tissues (B, D, F) of pigs inoculated with (A, B) H3N2 cold passaged mutant 531 [cp531] or H3N2 parental virus [par531], (C, D) H1pdmN2 cold passaged mutant 541 [cp541] or H1pdmN2 parental virus [par541], and (E, F) H1avN1 cold passaged mutants 1670 ST [cp1670ST] and 1670 MDCK-II [cp1670MDCK-II] or H1avN1 parental virus [par1670], respectively. *-*** indicate significance levels (p<0.05; <0.01; <0.001)

For both cp531 and its parent virus par531, viral antigen was detected in varying amounts in the lung, trachea, and nose (supplemental figure 6A). For cp531, scores given for lung and trachea were slightly higher (max score 3) than observed for par531 (max score 2). In the nasal mucosa, viral antigen was detected in the cp531 (max score 1) and par531 group (max score 3). No viral antigen was present in cp531 contact animals (not shown). In line with immunohistochemistry, histopathological changes of varying severity were present irrespective of the virus strain used as indicated in supplemental figure 6B. Necrotizing inflammation of bronchi and bronchioles was found in all animals inoculated with cp531 (max score 2.5) and par531 (max score 2). Suppurative inflammation was significantly more frequently found in cp531 pigs (max score 2), than in the par531 group (max score 1.5). Likewise, peribronchiolar lymphocytic cuffing associated with inflammation was consistently present in the cp531 group (max score 2) while comparable lesions were found in only two out of four par531 inoculated animals (max score 2). Inflammation of the alveolar septa was present in both groups (max score 2) whereas mild tracheal and nasal lesions were only sporadically seen. In contrast to cp531 and par531, the two cp531 contact animals showed only mild histopathological alterations.

As for the cp531 virus, eight pigs were intranasally inoculated with 10^6^ TCID_50_ of cp541. Results were compared to a group of eight age-matched pigs which received the same dose of par541. Inoculation-related clinical signs including increase in body temperature were not observed in any pigs inoculated with par541 or cp541 (supplemental figure 5C, D). Cp541 inoculated animals nasally excreted substantial viral loads although significantly reduced compared to par541 (Figure 5C), which is corroborated by viral loads measured in tissues of nasal conchae at day 4 (Figure 5D). Similarly, significantly reduced viral loads for cp541 were observed in respiratory tissues, particularly in deeper respiratory tissues (bronchi, BALF and lung lobes and tributary lymph nodes (Figure 5D). No seroconversion occurred within 4 days post infection.

In line with virological data, less viral antigen was detectable in the tissues of cp541 inoculated animals compared to the par541 group (supplemental figure 6C; exemplarily shown in Figure 6). While the lungs of five of eight animals inoculated with par541 were antigen-positive (max score 2), viral antigen was found in only one cp541 inoculated pig (score 1). Significantly higher antigen levels were also found in the trachea of par541 inoculated animals (max score 2) than in the cp541 group. Nasal viral antigen amounts were comparable in both groups (max score 3 and 2, respectively). Histopathologically, only minimal changes were detected in cp541 inoculated animals compared to its parental strain (supplemental figure 6D). While the majority of animals inoculated with par541 showed necrotizing (max score 3) and suppurative pulmonary inflammation (max score 2.5), only one cp541 pig revealed mild suppurative lesions (score 1). In contrast to the small number of affected cp541 inoculated animals, the par541 group showed significant lymphocytic peribronchial reaction (max score 3) and inflammation of the alveolar septa (max score 3). Mainly mild histopathological changes of the trachea were present in the par541 (max score 1) and cp541 groups (max score 2). Infection-related lesions were present in the nose of both groups (max score 3).

**Figure 6.**
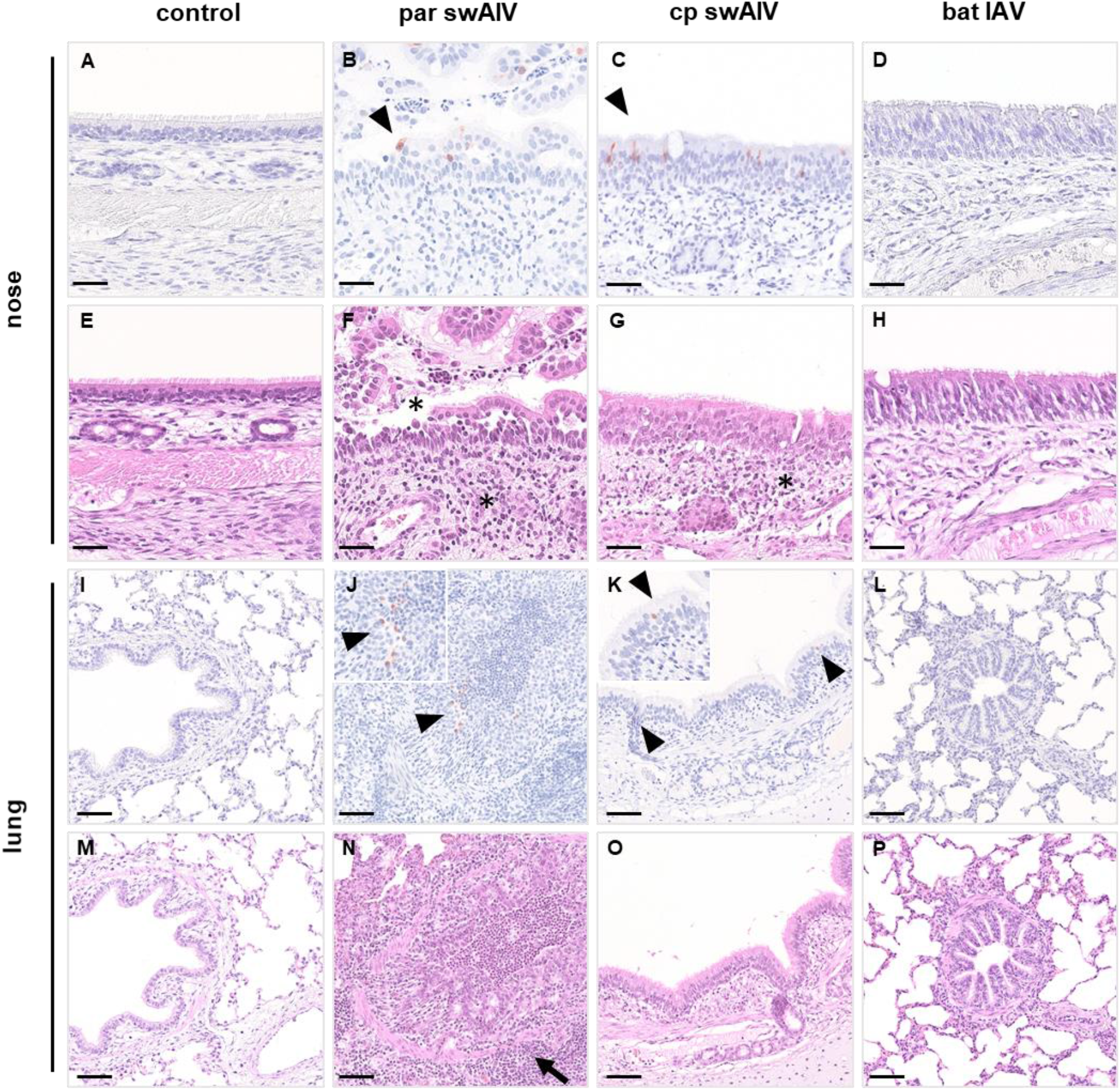
Representative histopathology and immunohistochemistry of the nose (A-D, IHC labeling; E-H, HE staining) and lung (I-L, IHC labeling; M-P HE staining) of control animals compared to pigs infected with par swIAV, cp swIAV or bat IAV. Antigen positive cells of the nose (B, C). bronchi and bronchioles (J-K) are indicated by arrowheads, insets demonstrate magnification of positive cells. Epithelial necrosis and immune cell infiltration of the nasal mucosa is shown by asterisk (F-G). Necrotizing bronchiolitis with peribronchiolar lymphocytic cuffing is highlighted by arrow (N). HE, Hematoxylin and eosin staining. IHC immunohistochemistry, Avidin-biotin-complex method, 3-amino-9-ethylcarbazole chromogen (red), hematoxylin counterstain (blue). Bars represent 50 µm (A-H) and 100µm (I-P).

For cp1670MDCK and cp1670ST, eight pigs each were intranasally inoculated with 10^6^ TCID_50_ of cp1670MDCK or cp1670ST. Results were compared to a group of eight age-matched pigs which received the same dose of par1670. Similar to the 541-virus pair, no clinical signs were observed (supplemental figure 5E-G). Interestingly, animals inoculated with the MDCK-II cp strain nasally excreted significantly less virus than those which received the ST-grown cp strain (Figure 5E). A trend for reduced replication efficacy of the cp1670MDCK-versus the ST-grown virus was also evident in deeper respiratory tissues (Figure 5F). Both cp viruses revealed significantly reduced replication compared to par1670 in the deeper parts of the respiratory tract. Seroconversion was not detected in any of the animals.

Consistent with the lower viral genome levels, all cp1670MDCK inoculated animals tested negative by immunohistochemistry (supplemental figure 6E). In contrast, inoculation with cp1670ST led to limited positive results only in the lung and trachea (max score 1), but achieved higher scores in the nose (max score 3) (exemplarily shown in Figure 6). In all animals inoculated with the par1670 strain varying amounts of viral antigen were detected in the lung (max score 3), trachea (max score 2) and nose (max score 3). Corresponding to immunohistochemistry, histopathological changes significantly differed between par1670, cp1670MDCK and cp1670ST (supplemental figure 6E, F). While pulmonary, tracheal and nasal lesions occurred frequently and were predominantly moderate to severe in par1670 infected pigs, lesions were mild and occurred only occasionally in the cp1670ST group or were completely absent in the cp1670MDCK group.

Respiratory clinical signs and fever in SPF pigs intranasally inoculated with swIAV are infrequent and dependent on different factors e.g. strain and dose, health status and age of the pigs. Therefore, and in line with the European Pharmacopoe used here as a guideline, virological (virus loads, excretion titers) and immunohistological rather than clinical readouts have been used to assess safety of swIAV vaccine candidates. On basis of this system, cp541 and cp1670 but not cp531 did reveal characteristics of an attenuated (*att*) phenotype, i.e. reduced viral replication in deeper respiratory tissues in the target species, pig.

### Ch-bt IAV replicated asymptomatically in pigs but ch-bt IAV H17-1670 replicated more extensively compared to H17-1670-NS1-2x

Eight pigs each were intranasally inoculated with 10^6^ TCID_50_ of H17-1670 or H17-1670-NS1-2x, and at day 1 p.i. two contact pigs were associated to each group. ch-bt IAV expressing H1av and N1av in the background of bt IAV H17 replicated in the inoculated pigs (Figure 7A, B). No inoculation-related clinical signs were registered in any of the pigs during the observation period (supplemental figure 7). All animals inoculated with H17-1670 excreted virus nasally on 2 and 4 dpi with substantial RNA loads (Figure 7A). In contrast, only two pigs inoculated with H17-1670-NS1-2x shed virus nasally, and the viral load was significantly lower compared to H17-1670 (Figure 8A). Nasal excretion of either chimera in contact animals was not detected, at least these animals remained PCR-negative in nasal swabs within the four-day observation period after initial contact. H17-1670 replicated to significantly higher loads than the NS1-2x-chimera in upper respiratory tissues (conchae, trachea), but was hardly detectable in lung tissue samples (Figure 7B). Sentinels of chimera H17-1670, although PCR-negative in nasal swabs, showed discrete virus presence in the bronchi as well as in BALF. Negligible replication in the examined tissue and swab samples was evident for chimera H17-1670-NS1-2x. No seroconversion of the animals was detectable within the four-day observation period.

**Figure 7.**
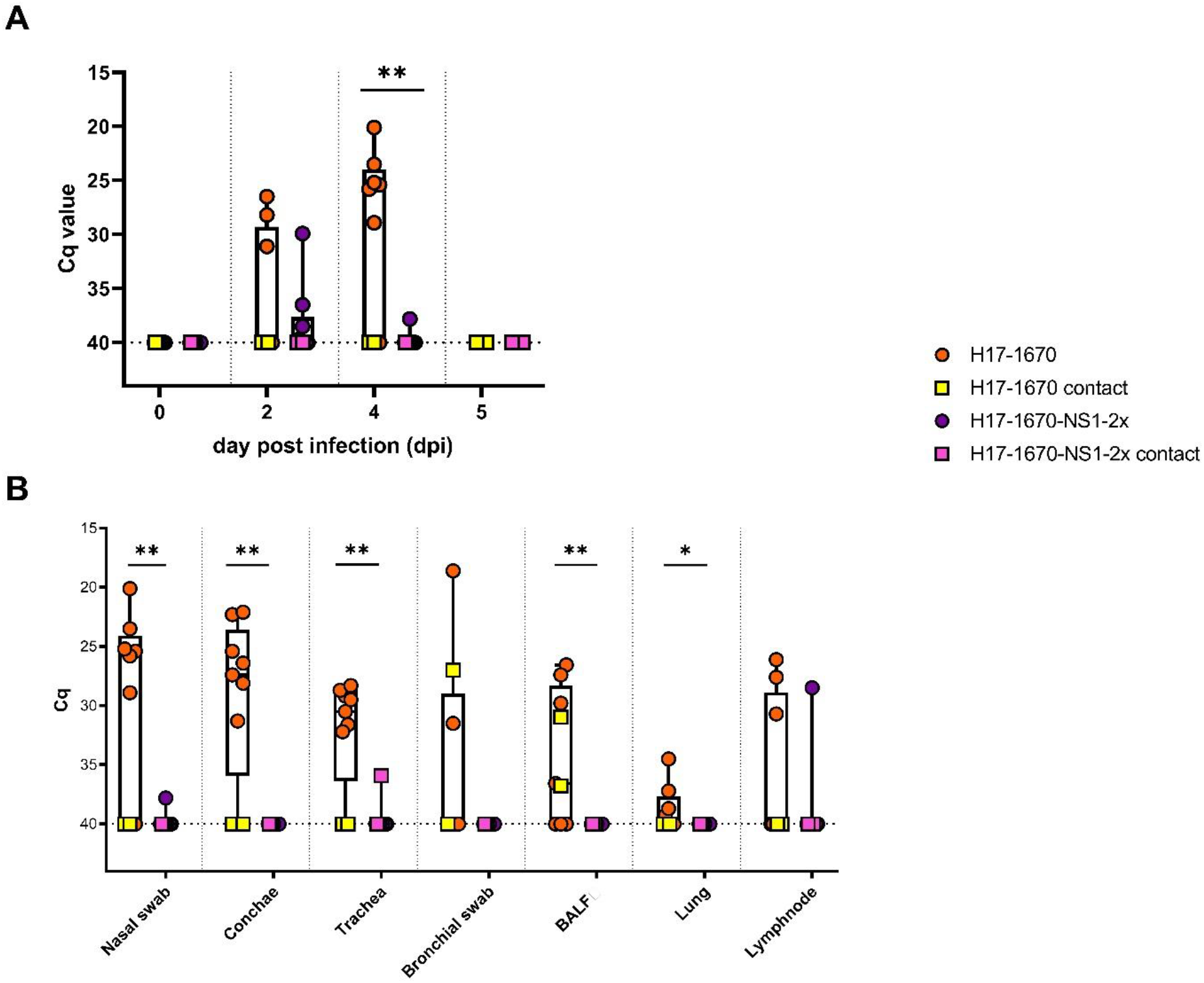
(A) Nasal viral excretion and (B) viral loads in respiratory tissues of pigs vaccinated with bat-swine Influenza A chimeric viruses [H17-1670] or [H17-1670-NS1-2x]. *-*** indicate significance levels (p<0.05; <0.01; <0.001)

**Figure 8.**
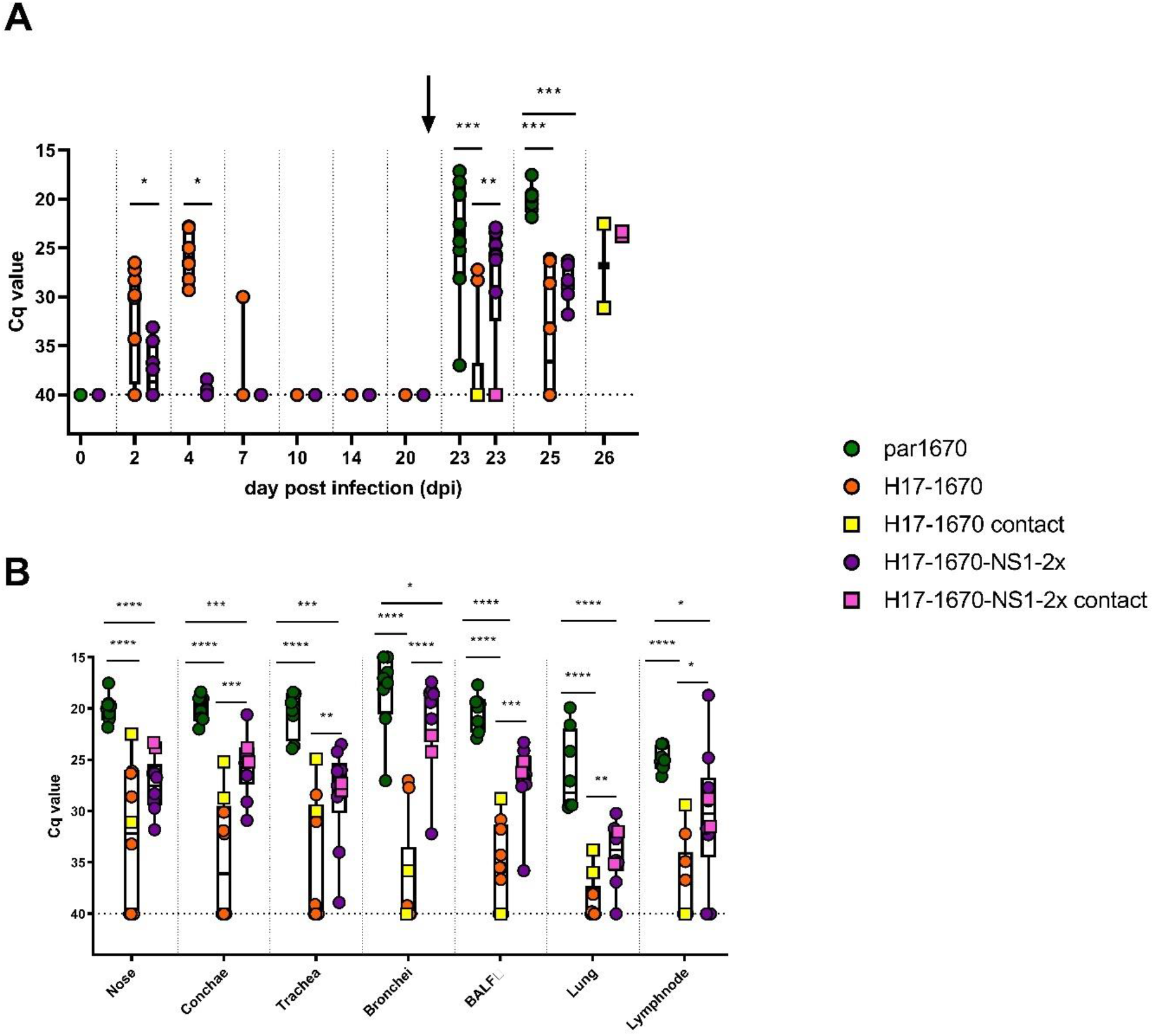
(A) Nasal viral excretion and (B) viral loads in respiratory tissues of pigs vaccinated with chimeric viruses [H17-1670] or [H17-1670-NS1-2x] and challenged with homologous virulent H1avN1 virus on day 21 post vaccination (black arrow). Green spheres show values of par1670 only for comparison; par1670 was not used for vaccination. *-**** indicate significance levels (p<0.05; <0.01; <0.001)

While, in summary, ch-bt IAV H17-1670 replicated efficiently in upper respiratory tracts and was also transmitted within four days of contact to sentinel pigs, H17-1670_NS1-2x seemed to be over-attenuated and incapable of efficient replication in the respiratory tract of pigs.

As presented in supplemental figure 8A, neither the vaccinated nor the contact animals showed positive immunohistochemistry. Mild histopathological changes in the lung and trachea were found only sporadically in individual animals (supplemental figure 8B).

### Ch-bt IAV H17-1670 conveys protective effects against challenge of vaccinated pigs

Due to the inconsistent safety studies of cp swIAV *in vitro* and in pigs in which cp531 did not clearly reveal an attenuated phenotype and because of the inconclusive mutation and reversion analysis of cp viruses, the focus concerning protective capacity was put on the ch-bt IAV. Eight pigs each were intranasally inoculated with 10^6^ TCID_50_ of H17-1670 or H17-1670-NS1-2x, and at day 1 p.i. each two contact pigs were associated. At day 21 p.i. the pigs were intranasally challenged with 10^6^ TCID_50_ of par1670. Each two naïve contact pigs were associated at day 1 post challenge. Animals vaccinated with chimera H17-1670 excreted virus at 2 and 4 dpi but not at 7, 10, 14 and 20 dpi. (Figure 8A). Detection of chimera H17-1670-NS1-2x in nasal swabs was limited to day 2 p.i. and negligible at 4 dpi with cq-values above 35 (Figure 8A).

After challenge with the homologous parental virus par1670 that has donated the HA and NA coding regions for the ch-bt IAV, at day 21 after vaccination no clinical signs of disease developed in either of the groups (supplemental figure 7). Animals of both vaccinated groups nasally excreted challenge virus at day 2 and 4 after challenge but at significantly lower levels than a group of eight mock-vaccinated pigs (Figure 8A, B). The mock-vaccinated group consisted of pigs that had been used for the safety studies of par1670. Significantly lower challenge virus loads in nasal swabs were found for animals of chimera H17-1670 compared to H17-1670-NS1-2x. Within four days of contact unvaccinated sentinels got infected and nasally shed virus on 5 dpc. A similar pattern as for nasal excretion was also evident for other respiratory tissues: RNA loads were lowest for H17-1670 vaccinated pigs and highest in mock-vaccinated animals while pigs vaccinated with H17-1670-NS1-2x took an intermediate position (Figure 8B).

In line with lower viral loads, also significantly less viral antigen was detected in all organs of the H17-1670 vaccinated group after challenge (supplemental figure 9A). While in H17-1670 vaccinated pigs substantial amounts of viral antigen were detected in the nose (max score 3), positive cells were only occasionally found in the lungs (max score 2) and were absent in the trachea (exemplarily shown in Figure 6). The lungs of the two contact animals of the H17-1670 challenge group were antigen-negative whereas the nasal mucosa of both animals and the trachea of one animal showed positive results. In contrast, the challenged H17-1670-NS1-2x group except for one inoculated animal, showed highly positive immunohistochemistry in all organs (max score 3). Likewise, the trachea and nose of both sentinels and the lung of one animal of the H17-1670-NS1-2x challenge group were antigen-positive. Overall histopathological alterations were most severe in par1670 inoculated animals (max score 3) while lesions, if present, were only mild to moderate in the two vaccinated groups (max score 2) (supplemental figure 9B). In particular, peribronchial lymphocytic cuffing and inflammation of the alveolar septa were significantly reduced in animals of the H17-1670 vaccinated group. Tracheal and nasal changes did not differ between the groups (max score 2 and 3, respectively). Histopathological alterations in contact animals of both vaccinated groups were mild to moderate or even absent.

In summary, despite strong attenuation, vaccination with chimera H17-1670-NS1-2x moderately reduced viral replication of the challenge virus par1670 compared to non-vaccinated animals, however, transmission to sentinels was not prevented. In contrast, vaccination with H17-1670 diminished viral replication of the par1670 to such an extent that transmission to sentinels was not observed, demonstrating the feasibility of bat chimeric viruses for live vaccine development.

## Discussion

Vaccination remains the most effective prophylactic measure to prevent influenza virus infection in pigs (Rahn, 2015; Sandbulte et al., 2015; Vincent, 2017). However, some of the virus strains used in the current vaccines in Europe were isolated more than 20 years ago. Although antigenic drift in swIAV seems not to be as striking as in human influenza strains uncertainty remains whether these vaccines can fully prevent formation of escape swIAV-variants. Recent studies have shown that resilient and broad protection against a wide range of antigenically diverse strains can be accomplished by serial boostering with different inactivated, adjuvanted vaccine strains; however, the type of strains and the order of booster vaccinations played a pivotal role in that study and had to be delineated empirically (Chepkwony et al., 2020). This leaves doubts regarding the practicability of such an approach in the field. Evidence from the field of the emergence of swIAV antigenic drift variants only recently raised questions about the use of the approved vaccines (Everett et al., 2019; Ryt-Hansen et al., 2020). This may be related to the altered epidemiology of the infection in large pig populations which can evidently favor a forced, population-specific antigen drift (Diaz et al., 2017; Rose et al., 2013; Ryt-Hansen et al., 2020). In face of the continuously increasing emergence of new swIAV variants in the field (Henritzi et al., 2020) and by considering the fact that the currently available vaccines on the European market solely comprise inactivated whole virus vaccines, novel concepts for swIAV vaccine design, ultimately aiming at a broader efficacy spectrum and widened flexibility for emerging strains, should be explored. Modified live-attenuated influenza vaccines (MLVs) mimic natural infection, are delivered intranasally, infect the upper respiratory tract and activate both humoral and cell-mediated immunity, thus providing more complete protection than inactivated vaccines (Sridhar, 2015). Several attempts to develop potentially more efficacious MLVs against swIAV infections in pigs have been reported previously (Loving et al., 2013; Pena et al., 2011). The potential benefits of MLVs must be weighed against the risks compromising biological safety (Martinez-Sobrido et al., 2018). Assessment of biological safety of MLVs includes the stability of the genetic markers linked to attenuation over several (at least five) passages in cell culture or in the target species and thus the exclusion of immediate reversion to the pathogenic phenotype. Furthermore, the (unintentional) horizontal spread of the MLV virus must be taken into account. An uncontrolled lateral spread from vaccinated animals could favor undesired mutations that affect antigenicity and/or attenuation. High-titred MLV virus excretion may also promote undesired transmission to other age or functional compartments of the herd or even to other species.

In the present study, two approaches to develop MLVs against swIAV were pursued, temperature-sensitive (*ts*) swIAV mutants and swIAV-bat IAV chimeras respectively: Our attempts to generate *ts* mutants of three antigenically diverse swIAV lineages by serial cold passaging of virus isolates in cell cultures did not yield conclusive results. Serial passaging in either MDCK or ST cell cultures under cold stress did not lead to significant mutations of HA and NA sequences, meaning the strains remained antigenically unaltered. Despite some growth-related indication of an adaptation of the three viruses to replication at 28°C, no previously described genetic markers of *ts* mutants of human, equine or avian influenza viruses could be identified (Broadbent et al., 2014; He et al., 2013; Meyer et al., 2016; Rodriguez et al., 2018; Townsend et al., 2001). Nevertheless, the majority of passage-associated mutations accumulating in the three strains during cold passaging occurred in genome segments of the polymerase complex, especially in the PB2 segment; which is in line with established *ts* mutants derived from human and equine IAV lineages. One mutation (E158G), which was independently selected in two out of four cold passaged viruses mightrepresent a marker. On the other hand, the plethora of coding mutations that differ between the four passage 60 viruses (Table 2) might signal ongoing selection and/or presence of multiple virus populations. Attempts to plaque-purify cp viruses failed (not shown), therefore bottleneck passaging at very low MOIs had to be applied. Presence of mixed and possibly instable virus populations was also suggested by the fast reversion of cp passage 60 strains to thermophilic and cytopathic replication at 37°C during serial blind passaging (supplemental table 1). The future way forward might be to introduce the D87G and E158G mutations selected here, and other, previously described *ts*-related mutations, into parental swIAV via reverse genetics. In our *in vivo* experiments in mice and pigs, at least two cp strains displayed an attenuated phenotype after nasal application. In particular, reduced replication in lung tissues but not in nasal conchae was observed compared to infections with parental viruses. Yet, pneumonic lesions of cp-infected mice at 10 dpi suggested lung-associated replication also of at least some of the cp isolates. Despite these promising biological features, the inconclusive genetic data of cp swIAV established here led us to focus further work on the second approach of generating MLVs against swIAV infections in pigs: ch-bt IAV.

It was shown by antigenic mapping that the expressed swIAV glycoproteins were antigenically authentic. Our studies showed that both viruses replicated in cell culture and in the target species pig, although the virus with additional attenuating mutations in the NS1 segment was considerably weaker in all tested systems (lower growth in MDCKII and ST cell cultures and highly significantly lower and shorter-lasting ability to replicate in mice and pigs). Despite moderately high viral loads that were found in nasal swabs of the animals in the chimera without the additional NS1-mutations, transmission of this chimera to two sentinel animals could not be clearly demonstrated. However, natural bat-IAV infections have not yet been associated with pathogenicity for humans or non-bat mammals or other species and there is currently no evidence that the HA/NA replacement in bat-associated IAVs of subtype H17N10 could significantly increase the risk potential of these viruses, provided that only HAs expressing a trypsin-sensitive cleavage site are used. In the end, our challenge experiments with vaccinated animals demonstrated a protective immunity which was mainly characterized by a highly negative challenge virus replication in the lower airways. The protective effect with regard to the measured viral loads was significantly lower in the group that received the NS1-attenuated chimera, yet still increased when compared to unvaccinated animals. A similar H17N10-based vaccine study was performed by the group of Lee et al. (2021). In this case, a H3N2 based bat-IAV chimera was generated expressing a truncated NS1 (NS aa1-128), designated MLV1. Vaccination with MLV1 protected against heterologous viral challenge and the truncation mutant replicated to high viral lung titer compared to H17-1670-NS1-2x-infected swine. While the two mutations in NS1 abrogate the RNA-binding activity of NS1, the ability to bind RNA is not affected in the truncated NS1 protein. Since comparable virus doses were used in both studies, it seems that the truncated NS1 variant is less attenuated than a NS1 variant with defective RNA binding domain. Of note, MLV1, H17-1670-NS1-2x and ch-bt IAV H17-1670 expressing an intact NS1 did not develop any disease.

These findings are therefore in line with former studies, where chickens and ferrets inoculated with bat-IAV chimeras expressing (monobasic) HA and NA of a highly pathogenic avian influenza virus of subtype H5N1 did not develop disease nor pathological alterations but were protected against homologous challenge. Chimeras instantly replicated to high titers in ferrets but required multi-passage adaptation in chickens (Ren et al., 2019; Schon et al., 2020). Furthermore, the unsurpassed efficacy of MLVs for induction of mucosal immunity has been shown for various IAV candidates and licensed vaccines in North America based on NS1 truncations or temperature-sensitive mutants (Genzow et al., 2018). However, due to reassortment events with wild-type IAV, indirect reversion to virulence occurred for the North American vaccine (Sharma et al., 2020). All previous attempts were lacking a bona fide mechanism that prevented reversion to virulence by blocking reassortment with co-circulating field viruses. Due to the genetic backbone of bat-IAV, bat-chimeric viruses cannot reassort with conventional influenza viruses from birds and non-bat mammals (Ciminski et al., 2017; Juozapaitis et al., 2014; Ma et al., 2015), thereby preventing reversion to virulence. This is in fact a safety feature of future MLVs based on such bat-IAV chimeras. On the other hand, MHC class II has recently been identified to act as a receptor for bat-IAV through interaction with its HA protein. The broad binding capacities to MHC class II of various mammalian species has raised concerns for its zoonotic potential (Karakus et al., 2019). However, in our studies, the potential factor for zoonotic susceptibility (the authentic bat-IAV HA), was replaced by the subtype H1av of swIAV origin with restricted binding preferences for sialic acid receptors.

These findings suggest that even if infection levels are significantly reduced by an antigenically matched vaccine, thereby slowing both the dissemination of the virus through a herd and the generation time for naive contact pigs, transmission to naive animals may not be entirely interrupted. These findings are in agreement with those from previous reports indicating that vaccines are able to significantly limit nasal shedding, irrespective of the vaccine platform used, may delay viral transmission to naive pigs in direct contact and may also decrease the likelihood of indirect transmission (Loving et al., 2013; Ryt-Hansen et al., 2019), presumably by lowering viral load in the environment.

Based on the apparently stable backbone of the chimeric candidates tested, HA and NA genome segments of further antigenically differentiable swIAV lines (H1huN2, HpdmN1, H3N2) could be generated by reverse genetics. Testing whether multi-(tetra)valent *ts*-MLV induces a balanced and robust protective immune response, similar to the licensed quadrivalent *ts* vaccine for humans, in safety and vaccination-challenge studies in pigs would be desirable.

## Material and Methods

### Virus origin and cell culture propagation

#### Parental swIAV

SwIAV parental (wild-type) strains were recruited from the study by Henritzi et al. (2020). Representative isolates of three virus lineages currently circulating in European domestic pig populations were selected. These included H1avN1 (A/swine/Germany/AR1670/2014, abbr. 1670), H1pdmN2 (A/swine/Germany/R541/2012, abbr. 541), and H3N2 (A/swine/Netherlands/AR531/2015, abbr. 531). These viruses were propagated in Madin-Darby canine kidney cells (Madin et al., 1958) (MDCK-II, FLI Collection of Cell Lines in Veterinary Medicine CCLV-RIE 1061) in Eagle’s Minimal Essential Medium (MEM) supplemented with 10,000 U/ml Penicillin-Streptomycin (Thermo Fisher Scientific, Germany) and 2 µg/ml L-1-Tosylamide-2-phenylethyl-chloromethyl-ketone (TPCK)-treated trypsin (Sigma-Aldrich, USA). Viral infectivity titers were measured in MDCK-II cells in 96-well plates, which were incubated for three to five days at 37°C in a 5% CO_2_ atmosphere. Induction of cytopathic effects (CPE) was examined natively. In addition, viral antigen was visualized using an immunoperoxidase monolayer assay (IPMA) as previously detailed (Postel et al., 2011). Briefly, cells were heat-fixated in 96-well plates and a sandwich assay with NP-specific monoclonal antibody H16-L10-4R5 (www.atcc.org) and a peroxidase-coupled goat anti-mouse IgG HRP polyclonal antibody (Bio-Rad Laboratories GmbH, Germany) was carried out. 3-amino-9-ethylcarbazol (3-AEC; Sigma Aldrich, Germany) was used as a chromogen. TCID_50_ titers were calculated according to Kärber (1931).

#### Virulent viruses used as control and for challenge experiments

The murine-adapted, virulent H1N1 strain A/Puerto Rico8/1934 [PR8] served as a positive control in mice inoculation experiments as it is known to induce clinically severe pneumonia with histopathological lesions in lungs of mice (Fukushi et al., 2011). For challenge of pigs, the (homologous) parental virus strain 1670 (H1avN1) was used.

#### Serial cold-passaging of swIAV

Sixty serial passages of the three selected wild-type viruses were generated in parallel in MDCK-II cells, as well as in the immortalized porcine cell line “swine testicle” (ST, FLI Collection of Cell Lines in Veterinary Medicine CCLV-RIE 0606), respectively. As the passages progressed, the incubation temperature was gradually lowered from 37°C to 33°C and finally to 28°C. Similarly, the inoculation dose was gradually reduced from 1 mL supernatant of the previous passage to 100 and 10 µL for a 25 cm^2^ cell culture flask. Figure 2 provides details of the passage history. The identity of HA and NA subtypes during serial cold passages (cp) was confirmed every 10 passages by subtype-specific tetraplex real-time PCRs [RT-qPCR] as described by Henritzi et al. (2016). To obtain information on the frequency of reversion to the thermophilic phenotype, the cp candidates were subsequently cultured ten times at 37°C in cell culture.

#### Bat IAV

Ch-bt IAV were generated by the eight-plasmid reverse-genetics system described by Juozapaitis et al. (2014). In short, eight pHW2000 plasmids encoding all viral segments were transfected into HEK 293T cells using Lipofectamine 2000 (Thermo Scientific). Plaque assays of viral rescue supernatants were performed on MDCK-II cells to generate single plaques used for further stock generation on MDCK-II cells, in case of the NS-wt virus, or MDCK-NS1-NPro cells, in case of the NS1-2x virus. TPCK-treated trypsin (1µg/ml) was added to viral growth medium to allow multicycle replication of the viruses. Correct introduction of the NS1 mutations was verified via cDNA synthesis of viral RNA using the One-Step RT-PCR kit (Qiagen, Hilden, Germany), amplification and sequencing using NS-specific primers. For viral growth kinetics, MDCK-II, ST, and MDCK-NS1-Npro cells were infected with the indicated viruses at a MOI of 0.001. Supernatant was harvested at the indicated timepoints and viral titers were determined via plaque assay.

### Molecular investigations

#### Construction of chimeric-bat IAV

Two reassortment-incompetent ch-bt IAV were generated as shown in Figure 2. The parental swIAV isolate A/swine/Germany/AR1670/2014 (H1avN1) served as donor of the HA and NA segments which were flanked by authentic 5’ and 3’ sequences of A/little yellow-shouldered bat/Guatemala/164/2009 (H17N10) as described by Juozapaitis et al. (2014). The resulting chimera, H17-1670, harbored six unmodified internal segments of the H17N10. The second chimera, H17-1670-NS1-2x, consisted of five unmodified internal segments of the H17N10 bt IAV. Its NS segment was mutated to express the amino acid substitutions R39A and K42A. These mutations are supposed to attenuate the original H17N10 bt-IAV due to structural changes in the RNA binding domain of the NS-1 protein (Turkington et al. 2015).

#### Growth curve analyses

Viral growth kinetics were compared in MDCK-II and ST cell cultures at 28 and 37°C. Triplicates of cell-free supernatants were collected at 0, 1, 2, 6, 10, 24, 48, and 72 hours post inoculation [hpi]. TCID_50_ values were determined as described above.

#### Real time RT-PCR (RT-qPCR)

Viral RNA was purified from clinical samples or cell culture supernatants with the QIAamp®Viral RNA Mini Kit (Qiagen, Hilden Germany) according to the manufacturer’s instructions. TaqMan based RT-qPCRs were implemented using the AG-Path-One Step RT-PCR Kit (Ambion). Cycling was performed on a Bio-Rad CFX96 real-time PCR detection system (BioRad, Munich, Germany). Thermocycling conditions for the generic Matrix (M)-specific and HA/NA subtype-specific tetraplex RT-qPCRs (especially subtype H1avN1) were as described previously (Henritzi et al., 2016; Spackman et al., 2002).

#### Conventional One-Step RT-PCR and Sanger sequencing

The Superscript-III one step RT-PCR Kit with Platinum Taq polymerase (Invitrogen, Germany) was used for conventional RT-PCRs for each of the eight segments. Primers and optimized thermocycling conditions on an Analytik Jena Flex Cycler for full genome segment RT-PCRs are available on request. Size-separated amplificates were purified from agarose gels using the QIAquick Gel Extraction Kit (Qiagen, Hilden, Germany). Purified PCR products were used directly for cycle sequencing reactions (BigDye Terminator v3.1 Cycle Sequencing Kit, Applied Biosystems) which were purified using NucleoSEQ columns (Macherey-Nagel, Germany), and sequenced on ABI PRISM 3100 and 3130 Genetic Analyzers (Life Technologies). Nucleotide sequences were curated and assembled using the Geneious software suite (Biomatters Ltd.) (Kearse et al., 2012).

#### Next and third generation sequencing – Illumina and MinION

RNA was amplified with influenza-specific primers (Hoffmann et al., 2001) using Invitrogen Superscript III One-Step RT-PCR with Platinum Taq (ThermoFisher Scientific, Waltham, USA). The simultaneous amplification of all influenza segments is based on a one-step RT-PCR method along with primers designed to bind to the conserved 3’ and 5’ ends of the segments. After amplification, purification of the PCR products was executed with AMPure XP Magnetic Beads (Beckman Coulter, Fullerton, USA). Quantification was finally conducted with the NanoDrop™ 1000 Spectrophotometer (ThermoFisher Scientific). The library preparation for the MinION platform was accomplished with the Rapid Barcoding Kit (RBK-004, Oxford Nanopore Technology, Oxford, UK) following the manufacturer’s instructions. Finally, mapping and consensus production of the quality checked and trimmed reads was conducted in the Geneious Software Suite (v11.1.5; Biomatters) with MiniMap2 (Li, 2018), creating full genomes for all samples. Illumina sequencing was conducted by the company Eurofins according to their in-house NGSelect DNA protocol with fragmentation, adapter ligation with UDIs (unique dual indexing) producing up to 5 million paired reads of 150 bp in length. Mapping and consensus genaration of the quality checked and trimmed reads was conducted in the Geneious Software Suite (v11.1.5; Biomatters) using Bowtie2.

### Experimental animal infections

#### Ethics statement

All animal experiments were approved by the State Office for Agriculture, Food Safety and Fishery in the Federal State of Mecklenburg-Western Pomerania, Germany: LALFF M-V 7221.3-1-030/19. All animals were kept under BSL-2 (cold-passaged swIAV) or BSL-3 (ch-bt IAV infections) conditions in the corresponding animal facilities at Friedrich-Loeffler-Institute (FLI), Germany.

All reverse genetically constructed chimeric viruses carrying gene segments of mammalian or avian influenza A viruses in the backbone of bat influenza A viruses were approved by the Regional Council of Baden-Württemberg, Germany; and the State Office for Health and Social Affairs of Mecklenburg-Western-Pomerania, Germany: SSI2-UNI.FRK.05.23/05.18/05.22 and LAGuS3021_4/11.5.17.

#### Animal origin

In order to control possible host-specific influences on the course of the infection, a total of 120 SPF C57BL/6 mice from two different breeding lines were tested (from in-house breeding at FLI (Greifswald-Riems, Germany) and from Charles River Laboratories breeding (Research Models and Services GmbH, Sulzfeld, Germany). Further, in total 102 six-week old German landrace weaner pigs aged 10 weeks were obtained from a commercial high health status, swIAV-negative herd (BHZP-Basiszuchtbetrieb, Garlitz-Langenheide, Germany). All purchased pigs tested negative in nasal swabs for swIAV RNA and IAV nucleocapsid protein-specific antibodies in serum by RT-qPCR and ELISA (ID Screen® Influenza A Antibody Competition Multi-species, IDVET, Germany), respectively. For the mouse infections with the ch-bt IAV vaccine candidates, 10 SPF C57BL/6 mice were acquired from Janvier (Strasbourg, France).

#### Experimental inoculation of mice

Groups of eight 5-13-weeks-old C57BL/6 mice were anesthetized with isoflurane and inoculated intranasally with 10^4^ TCID_50_ of cp swIAV or their corresponding parental (par) viruses in a volume of 40 µl (Deeg et al., 2017). One day post infection (dpi), four non-inoculated contact animals were co-housed. For 10 days, the body weights were measured daily and clinical observations recorded. Two days after infection, two inoculated animals were removed for histopathologic evaluation and antigen labeling using immunohistochemistry (IHC) in the lung and determination of viral loads in the nose and lung by RT-qPCR. All remaining animals were sacrificed at 10 dpi and viral loads were measured. Based on RT-qPCR results (cq value < 30), selected animals were examined for lung lesions and IHC was performed. For the infections with the ch-bt IAV, mice were anesthetized with a ketamine (100 µg per g of body weight) and xylazine (5 µg per g of body weight) mixture. Inoculation was performed intranasally with 10^4^ PFU in 40 µl PBS. The mice were sacrificed 3 dpi to harvest organs. Viral titers in lung and snouts of the infected mice were determined via plaque assay.

#### Experimental inoculation of pigs

##### Safety studies

The clinical sequelae of experimental infections with either cp swIAV, ch-bt IAV or par swIAV were studied. Study designs were related to the specifications of the European Pharmacopoeia (Ph. Eur., Monograph on Porcine Influenza Vaccine (inactivated); 01/2017:0963) where possible.

Eight pigs per virus were intranasally inoculated with 106 TCID_50_ of the respective viruses in a volume of 2 ml (1 ml per nostril) using mucosal atomization devices [MAD] (Wolfe Tory Medical, USA). Two non-inoculated contact animals were associated at 1 dpi. At 4 dpi, all inoculated pigs were sacrificed for dissection of the respiratory tract. One day later (4 days post contact, dpc), the contact animals were autopsied. Clinical scores (according to parameters detailed in the appendix) and body temperature measurements were recorded daily. Nasal swabs were obtained at days 0, 2, and post mortem, respectively. During post mortems, nasal conchae, trachea, a bronchial swab (left lung), a bronchoalveolar lavage (right lung), lung tissues from seven locations, and tracheobronchial lymph node were recovered for examination of virus load by RT-qPCR and histopathological changes based on the protocol by Gauger et al. (2012) and immunohistochemical investigations (for details see appendix, supplemental methodology). In addition, blood samples were collected at day 0 and post mortem.

##### Challenge studies

Protective effects of mucosal administration of ch-bt IAV were studied in challenge experiments commencing 21 days post inoculation (vaccination). Similarly to the procedures of the safety studies, vaccinated animals received 10^6^ TCID_50_ of the respective challenge viruses in a volume of 2 ml (1 ml per nostril) by MAD. On day 1 post challenge (dpch), each two non-inoculated contact animals were added to control challenge virus transmission. Nasal swabs were obtained at days 0, 2, and post mortem, respectively. At dpch 4 and 5, respectively, challenged and contact pigs were sacrificed and examined as described above.

### Histopathology and immunohistochemistry

Full autopsy was performed on all animals under BSL2 (cp viruses) or BSL3 (ch-bt IAV) conditions. For pigs, the lung, trachea and nose, for mice, the whole lung was collected and fixed in 10% neutral-buffered formalin. After trimming and paraffin-embedding, 2-3-μm-thick sections were stained with hematoxylin and eosin (HE). Consecutive slides were processed for immunohistochemistry according to standardized procedures of avidin-biotin-peroxidase complex-method as described (Schwaiger et al., 2019). The primary antibody against the IAV was applied overnight at 4°C (ATCC clone HB-64, 1:200 for pig tissues; rabbit anti-nucleoprotein serum 1∶750 for mice tissues), the secondary biotinylated goat anti-mouse or anti-rabbit antibody was applied for 30 minutes at room temperature (Vector Laboratories, Burlingame, CA, USA, 1:200). Slides were scanned using Hamamatsu S60 scanner, evaluation was performed using NDPview.2 plus software (Version 2.8.24, Hamamatsu Photonics, K.K. Japan). Evaluation and interpretation was following post examination masking approach (Meyerholz et al., 2018).

For mice, HE stained whole lung slides were evaluated for perivascular, peribronchiolar, and alveolar immune cell infiltrates as well as pneumonia-associated atelectasis, bronchiolar epithelial necrosis and regenerative hypertrophy/hyperplasia. All changes were recorded on ordinal scores using the tiers: 0 = no changes, or 1 = focal to oligofocal (<5% of the lung affected), 2 = multifocal (6-40%), 3= coalescing (41-80%), 4 = diffuse (>81%) changes. The sum of all values resulted in the pneumonia score. The distribution of viral antigen, four lung lobes were semi-quantitatively scored on an ordinal 0 to 4 scale: 0 = negative; 1 = focal or oligofocal, 2 = multifocal, 3 = coalescing, and 4 = diffuse immunoreactive bronchiolar and alveolar epithelial cells, respectively. The sum of all values resulted in the antigen score.

For pigs, standardized histopathological evaluation of all lung lobes and tracheal sections was performed based on the protocol published by Gauger and colleagues (Gauger et al., 2012). In addition, viral antigen-associated nasal lesions were scored on a scale from 0 to 3 (score 1 = epithelial necrosis only; score 2 = epithelial necrosis and oligofocal (≤4) rhinitis; score 3 = epithelial necrosis and multifocal (>4) rhinitis). Positively stained sections were semiquantitatively investigated for the amount and distribution of viral antigen. The most affected area per organ (lung, trachea and nose) was selected and the number of positive cells was counted in 40x (0.1 mm^2^). Subsequently, the distribution pattern of positive areas of the section was determined with 1= focal (n=1), 2= oligofocal (n=2-4), 3= multifocal (n>4). The number of positive cells was then multiplied by the distribution score, which gave the final value. The following final values were assigned the respective scores in lung: score 1=1-75; score 2= 76-150, score 3 >150. The scores in trachea and nose were assigned as follows: score 1= 1-20, score 2= 21-40, score 3= >40. For subsequent analyses, the max values of all scores per organ were included.

#### Statistics and illustration

The two-tailed Mann-Whitney-test was used to calculate significance of differences between trial groups; p <0.05 was considered significant. Graphical illustrations were produced using GraphPad Prism version 9.0.0 for Windows (GraphPad Software, La Jolla, CA). Histopathology figures were created using Adobe Photoshop version CS5 (Adobe Systems Software Ireland Limited, Ireland).

## Acknowledgments

The authors thank Aline Maksimov, Diana Parlow, Silvia Schuparis and Gabriele Czerwinski for outstanding technical assistance. For excellent care of animals and support during trials, we thank the animal keepers Steffen Kiepert, Christian Lipinski, Lukas Steinke, David Truschinski, Kerstin Kerstel, Frank Klipp, Doreen Fiedler, Ilona Bauernschäfer and necropsy assistants Ralf Redmer, Ralf Henkel and Christian Loth.

## Author Contributions

Conceived and designed experiments: TCH, MB, MS, FD, AG, PPP, DH. Acquired animal samples: AG, TCH, PPP, JSE, AB. Sample processing: AG, PPP, JSE, DH, AB, JK, AP. Pathological and histological investigation: JSE, AB. Data analysis, statistics, figure design: AG, TCH, PPP, MS, JSE, DH, AB, JK, AP. Manuscript preparation: AG, TCH, JSE, AB, PPP, MS. All authors reviewed and approved the final version of the manuscript.

## Data availability

All data generated or analysed during this study are included in the manuscript and supporting files.

