## Supplemental material for "Cold-passaged isolates and bat-swine influenza A chimeric viruses as modified live-attenuated vaccines against influenza A viruses in pigs"

**Supplemental table 1.** Reversion of cold passaged swIAV (passage 60) to cytopathic thermophilic replication in serial blind passages in cell culture at 37°C.

* - no CPE; + CPE detectable

| ID virus strain | Subtype | MDCKII | | | | | | | | | | ST | | | | | | | | | |
| --- | --- | --- | --- | --- | --- | --- | --- | --- | --- | --- | --- | --- | --- | --- | --- | --- | --- | --- | --- | --- | --- |
|  |  | 1 | 2 | 3 | 4 | 5 | 6 | 7 | 8 | 9 | 10 | 1 | 2 | 3 | 4 | 5 | 6 | 7 | 8 | 9 | 10 |
| A/swine/Netherlands/  AR531/2015 | H3N2 | - | - | - | - | - | - | - | - | - | + | - | - | - | - | - | - | - | - | - | - |
| A/swine/Germany/  R541/2012 | H1pdmN2 | + | + | + | + | + | + | + | + | + | + | - | - | - | + | + | + | + | + | + | + |
| A/swine/Germany/  AR1670/2014 | H1avN1av | - | - | - | - | - | - | - | + | + | + | - | - | - | - | - | - | - | + | + | + |

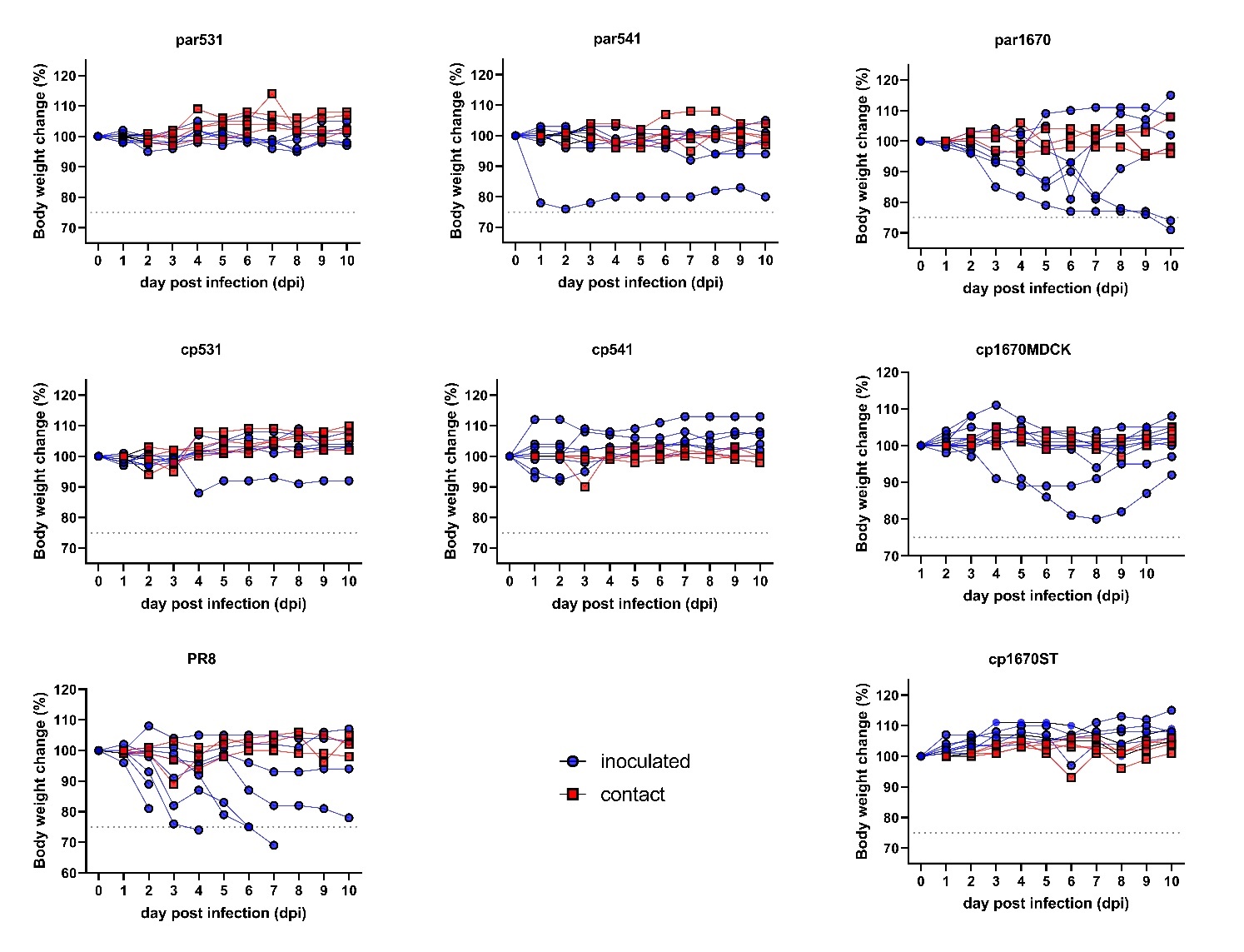

**Supplemental figure 1.** Kinetics of body weights of C57BL/6 mice inoculated intranasally each with 40 µL of parental (par) or cold-passaged (cp) strains 531 (H3N2, A, B), 541 (H1pdmN2; C, D) and 1670 (H1avN1, E, F, G) of swAIV, or with the mouse-adapted A/Puerto Rico/8/1934 (PR8) virus (H) used here as positive control. A weight loss of 25% was used as a humane endpoint.

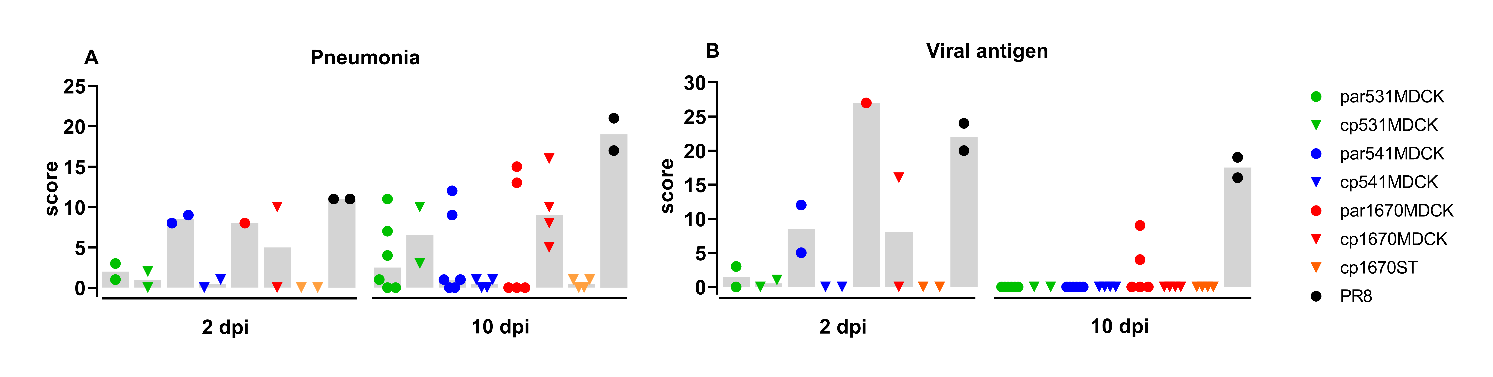

**Supplemental figure 2.** Pneumonia and viral antigen scores in the lungs of swIAV or PR8 infected C57BL/6 mice, 2 and 10 dpi. (A) Pneumonia score. The sum score for perivascular, peribronchiolar, and alveolar immune cell infiltrates, atelectasis, bronchiolar epithelial necrosis and regenerative hypertrophy/hyperplasia: score 0 = no changes, or 1 = focal to oligofocal (<5%), 2 = multifocal (6-40%), 3= coalescing (41-80%), 4 = diffuse (>81%) changes. Shown is the sum of all values recorded on the whole lung slides (maximum score = 24). (B) Viral antigen score. Viral antigen was semi-quantitatively scored on 4 lung lobes per animal: 0 = negative; 1 = focal or oligofocal, 2 = multifocal, 3 = coalescing, and 4 = diffuse immunoreactive bronchiolar and alveolar epithelial cells, respectively. Shown is the sum of all values per animal (maximum score 32). Bars refer to median.

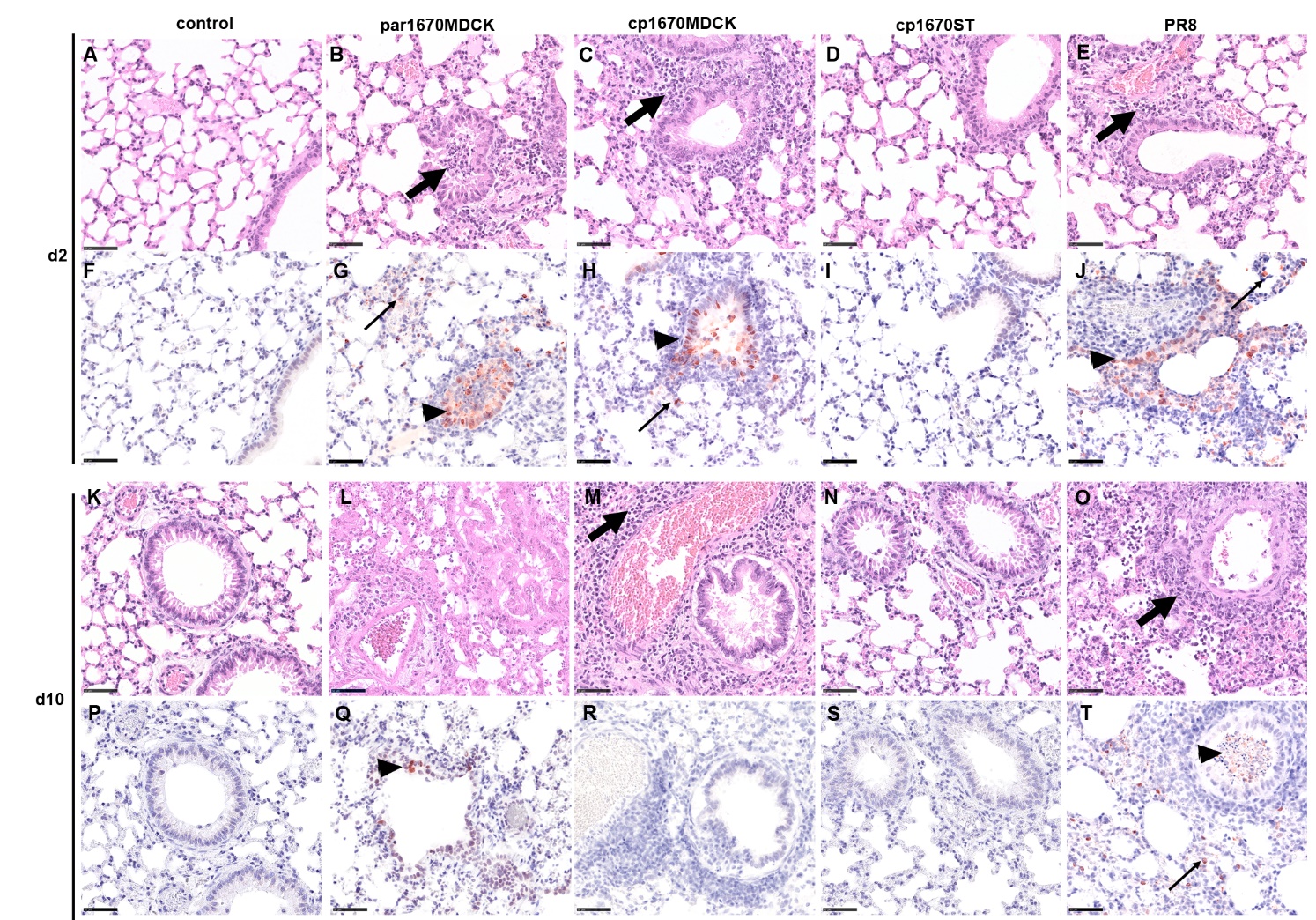
**Supplemental figure 3**. Representative histopathologic changes (A-E, K-O, HE staining) and lesion-associated viral antigen labeling (F-J, P-T, IHC labeling) in the lungs of swIAV (par or cp1670) or PR8 infected C57BL/6 mice, 2 (A-J) and 10 (K-T) dpi. Bronchiolar necrosis and immune cell infiltrates (bold arrows) and viral antigen in bronchiolar cell or intraluminal debris (arrow head) and alveolar epithelium (slender arrow). HE hematoxylin and eosin staining. IHC immunohistochemistry, Avidin-biotin-complex method, 3-amino-9-ethylcarbazole chromogen (red), hematoxylin counterstain (blue). Bars = 50 µm

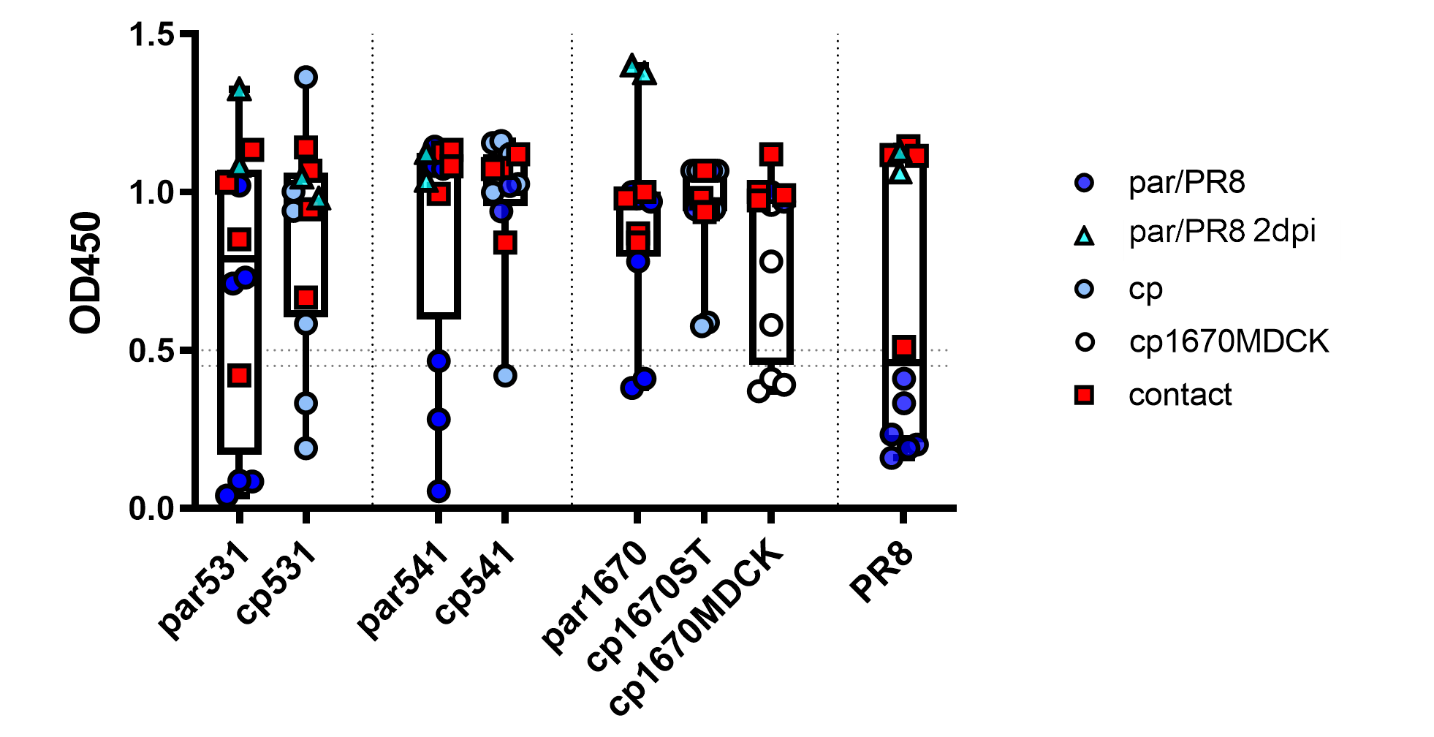

**Supplemental figure 4.** Development of nucleoprotein-specific antibodies against influenza A virus nucleoprotein as measured by competitive ELISA (ID-VET NP) at days 2 and 10 post infection. C57BL/6 mice were inoculated intranasally each with 40 µL of parental (par) or cold-passaged (cp) strains of swAIV, or with the mouse-adapted A/Puerto Rico/8/1934 (PR8) virus used here as positive control. OD value < 0.45 are considered positive.

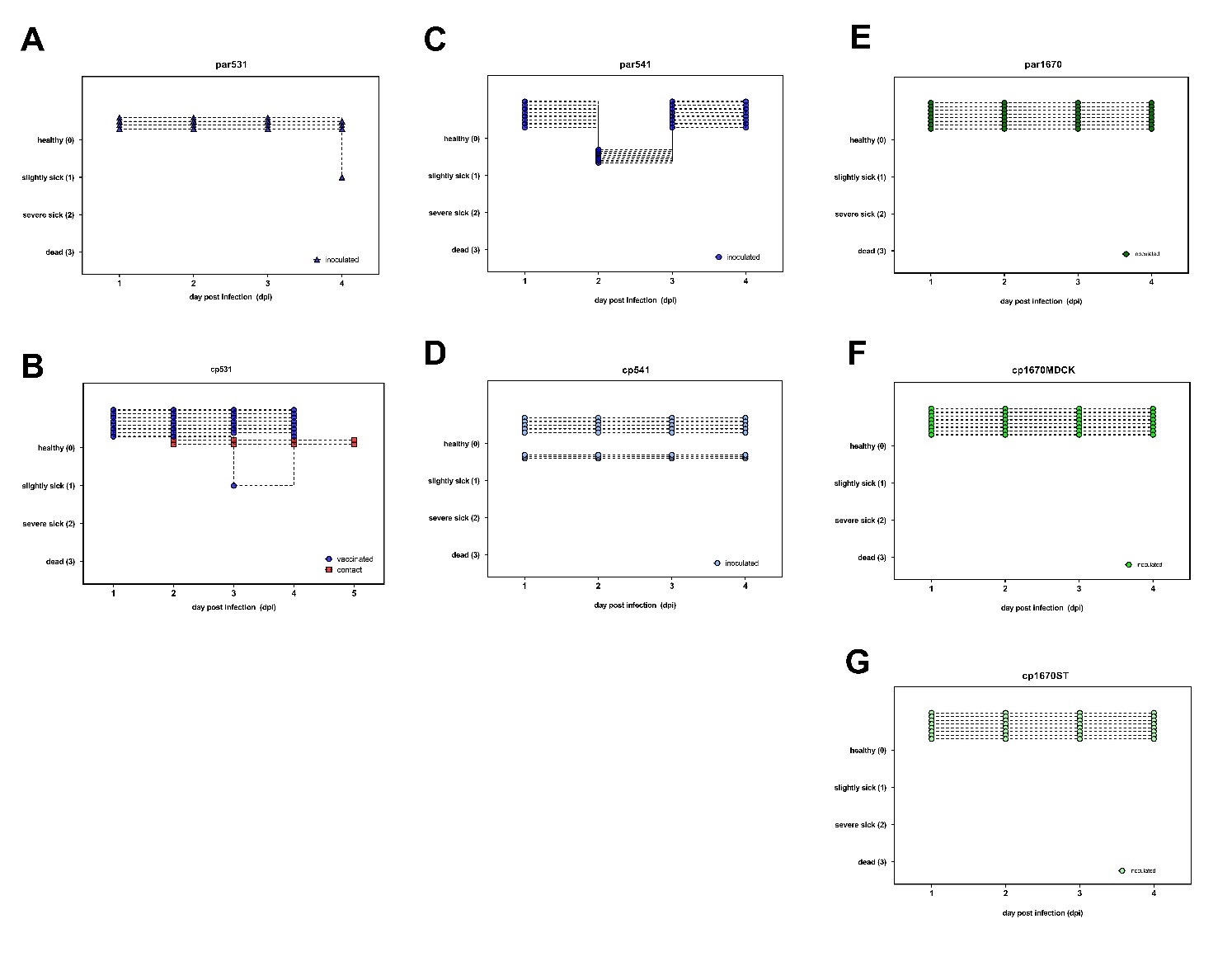

**Supplemental figure 5.** Clinical scores of pigs intranasally inoculated 10^6^ TCID_50_ of parental (par) and cold-passaged (cp) swIAV strains 531 (H3N2, A, B), 541 (H1pdmN2; C, D) and 1670 (H1avN1, E, F, G). Two contact pigs had been associated at day 1 post inoculation. Animals were kept for four days post inoculation or contact, respectively.

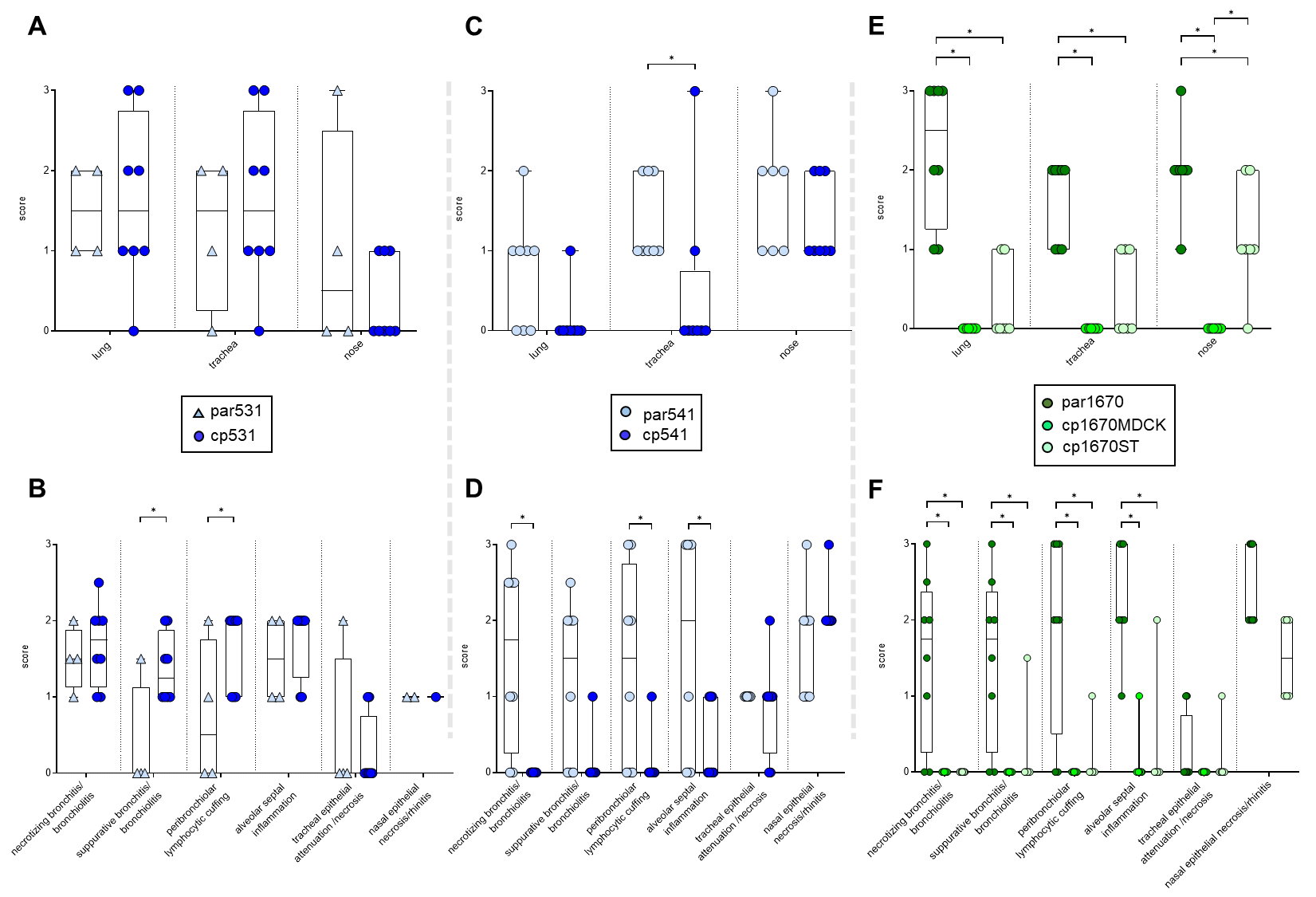

**Supplemental figure 6**. Immunhistochemical (upper panels) and histopathological (lower panels) semiquantitated alterations in tissues of each eight pigs intranasally inoculated with 10^6^ TCID_50_ of parental (par) or cold-passaged (cp) swIAV strains 531 (H3N2, A, B), 541 (H1pdmN2; C, D) or 1670 (H1avN1, E, F). Animals were sacrificed for pathological investigations at four days post inoculation. Significance levels are indicated by asterisk (*) (p<0.05).

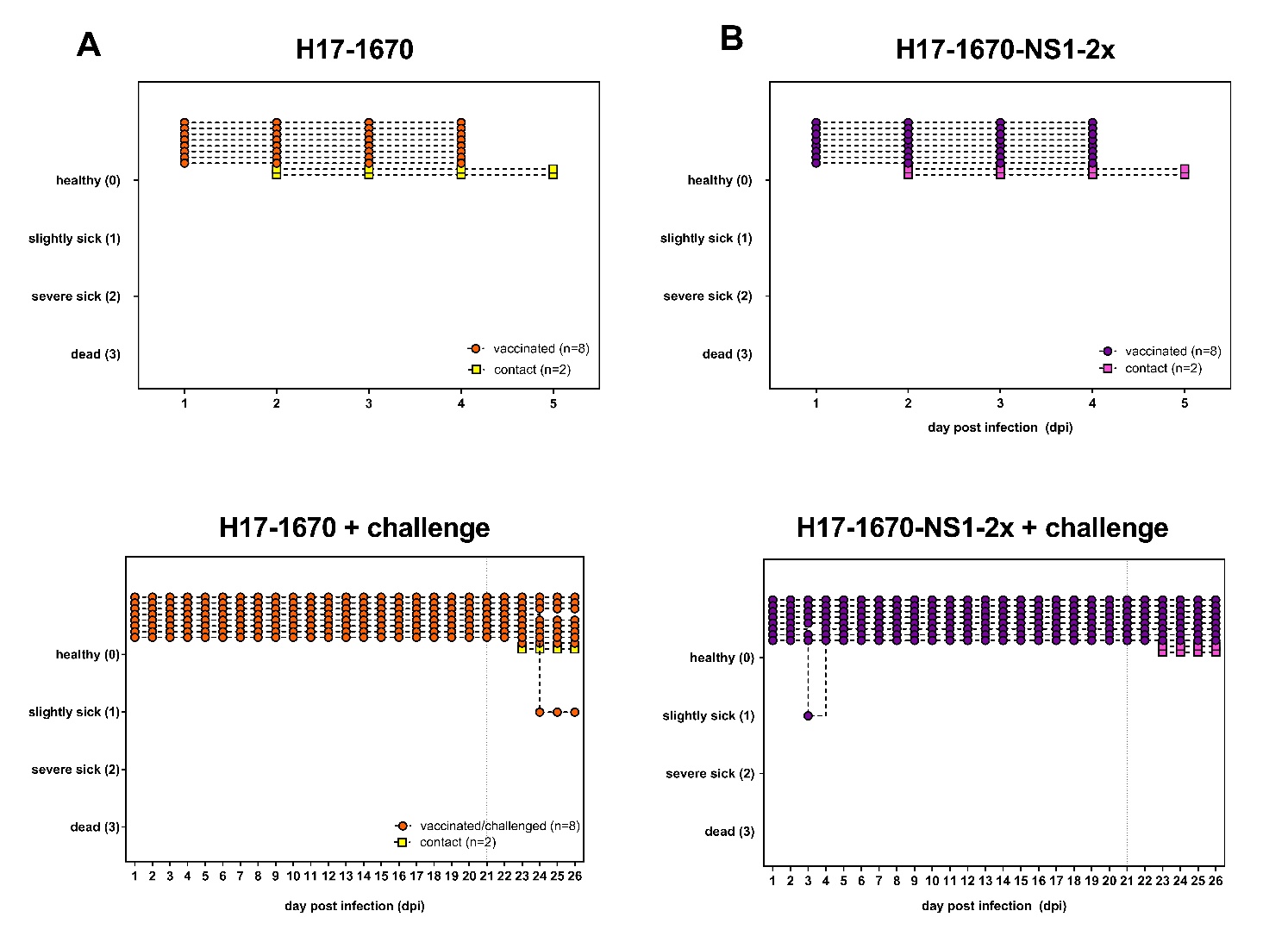

**Supplemental figure 7**. Clinical scores of pigs intranasally inoculated 10^6^ TCID_50_ of chimeric bat influenza A viruses H17-1670 (A) or H17-1670-NS1-2x (B) expressing hemagglutinin and neuraminidase proteins of swIAV 1670. Two contact pigs had been associated at day 1 post inoculation. Upper panels: Animals were sacrificed after four days post inoculation or contact, respectively, for pathological investigations. Lower panels: Animals were kept for 21 days and then challenged using homologous parental swIAV 1670.

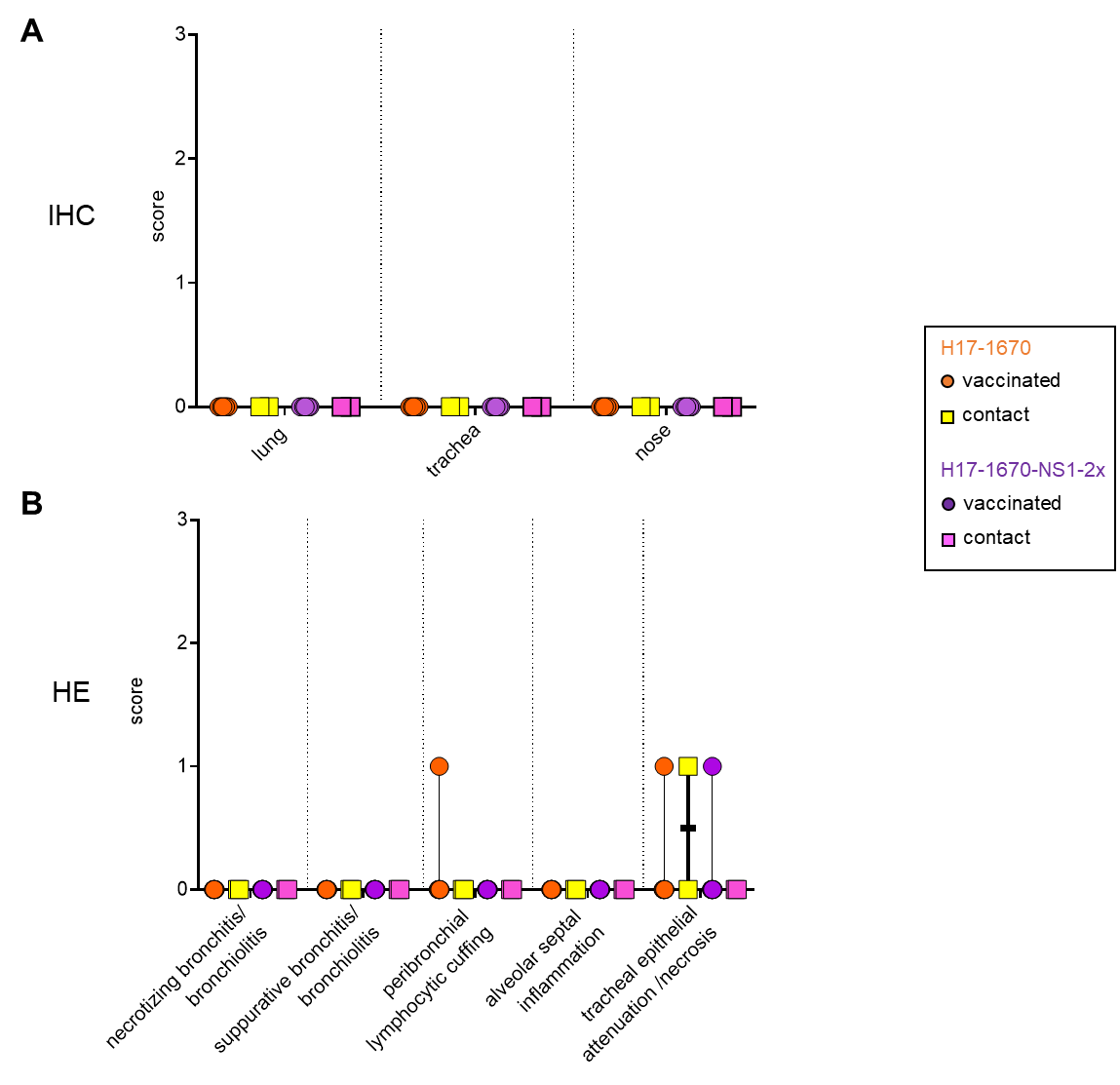

**Supplemental figure 8**. Immunohistochemical (A) and histopathological (B) semiquantitated alterations in tissues of each eight pigs intranasally inoculated with 10^6^ TCID_50_ of chimeric bat influenza A viruses H17-1670 or H17-1670-NS1-2x expressing hemagglutinin and neuraminidase proteins of swIAV 1670. Two contact pigs had been associated at day 1 post inoculation. Animals were sacrificed for pathological investigations at four days post inoculation or contact, respectively.

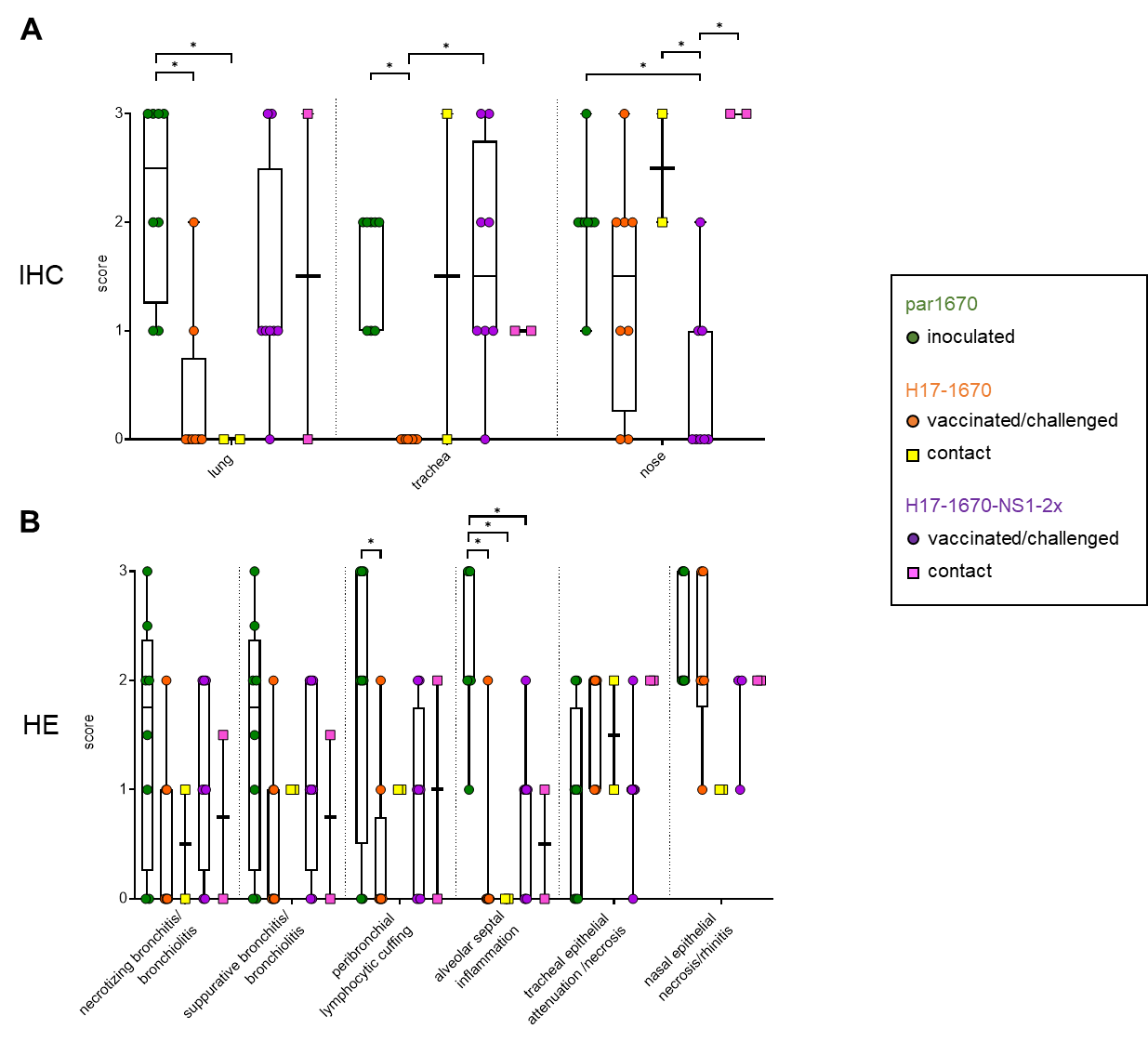

**Supplemental figure 9**. Immunohistochemical (A) and histopathological (B) semiquantitated alterations in tissues of each eight pigs intranasally inoculated with 10^6^ TCID_50_ of chimeric bat influenza A viruses H17-1670 or H17-1670-NS1-2x expressing hemagglutinin and neuraminidase proteins of swIAV 1670. Animals were challenged using homologous parental swIAV 1670 at day 21 after inoculation. Two contact pigs had been associated at day 1 post challenge. Animals were sacrificed for pathological investigations at four days post inoculation or contact, respectively. Significant differences are indicated by asterisk (*) (p<0.05).
